# Differential UBE2H-CTLH E2-E3 ubiquitylation modules regulate erythroid maturation

**DOI:** 10.1101/2022.01.18.476717

**Authors:** Dawafuti Sherpa, Judith Müller, Özge Karayel, Jakub Chrustowicz, Peng Xu, Karthik V. Gottemukkala, Christine Baumann, Annette Gross, Oliver Czarnezki, Wei Zhang, Jun Gu, Johan Nilvebrant, Mitchell J. Weiss, Sachdev S. Sidhu, Peter J. Murray, Matthias Mann, Brenda A. Schulman, Arno F. Alpiand

**Author notes:** Corresponding author Correspondence (AFA). Institute of Diabetes and Regeneration Research, Helmholtz Centre Munich, Neuherberg, Germany (Oliver Czarnezki). Department of Molecular and Cellular Biology, College of Biological Sciences, University of Guelph, Guelph, Canada (Wei Zhang). These authors contributed equally (Dawafuti Sherpa, Judith Müller, Özge Karayel).

## Abstract

The development of haematopoietic stem cells into mature erythrocytes – erythropoiesis – is a controlled process characterized by cellular reorganisation and drastic reshaping of the proteome landscape. Failure of ordered erythropoiesis is associated with anaemias and haematological malignancies. Although the ubiquitin (UB) system is a known crucial post-translational regulator in erythropoiesis, how the erythrocyte is reshaped by the UB system is poorly understood. By measuring the proteomic landscape of *in vitro* human erythropoiesis models, we found dynamic differential expression of subunits of the CTLH E3 ubiquitin ligase complex that formed distinct maturation stage-dependent assemblies of structurally homologous RANBP9-and RANBP10-CTLH complexes. Moreover, protein abundance of CTLH’s cognate E2-conjugating enzyme UBE2H increased during terminal differentiation, which depended on catalytically active CTLH E3 complexes. CRISPR-Cas9 mediated inactivation of all CTLH E3 assemblies by targeting the catalytic subunit *MAEA,* or *UBE2H*, triggered spontaneous and accelerated maturation of erythroid progenitor cells including increased heme and haemoglobin synthesis. Thus, the orderly progression of human erythropoiesis is controlled by the assembly of distinct UBE2H-CTLH modules functioning at different developmental stages.

## INTRODUCTION

Cellular differentiation in multicellular organisms is often accompanied by programmed proteome reshaping and cellular reorganisation to accomplish cell-type specific functions. For instance, during myogenesis proliferative myoblasts undergo a differentiation programme with induction of specialised cytoskeletal proteins to form myofibrils in terminally differentiated myofibers (Chal & Pourquie, 2017; Le Bihan *et al*, 2015), whereas adipose stem cells induce differentiation cues controlling expression of proteins involved in lipid storage and lipid synthesis (Tsuji *et al*, 2014). Recently, global temporal proteomic analysis during neurogenesis of human embryonic stem cells revealed large-scale proteome and organelle remodelling via selective autophagy (Ordureau *et al*, 2021). A striking example of proteome remodelling is mammalian erythropoiesis, which is required for the generation of disc-shaped enucleated erythrocytes, whose unique topology dictates function of efficient red blood cell movement through the vasculature (Figure 1A). After several specialized cell divisions, erythroid progenitors progress through morphologically distinct differentiation stages known as pro-erythroblasts (ProE), early and late basophilic erythroblasts (EBaso and LBaso, respectively), polychromatic erythroblasts (Poly), and orthochromatic erythroblasts (Ortho), a process associated with erythroid-specific gene expression (Cantor & Orkin, 2002; Cross & Enver, 1997; Perkins *et al*, 1995; Pevny *et al*, 1991; Shivdasani *et al*, 1995), reduction of cell volume (Dolznig *et al*, 1995), chromatin condensation (Zhao *et al*, 2016), and haemoglobinization. Ejection of the nucleus at the reticulocyte stage (Keerthivasan *et al*, 2011) is followed by the elimination of all remaining organelles such as Golgi, mitochondria, endoplasmic reticulum, peroxisomes, and ribosomes (Moras *et al*, 2017; Nguyen *et al*, 2017). The progression of erythroid maturation must be tightly controlled, although the molecular regulation of this process is not fully understood.

**Figure 1.**
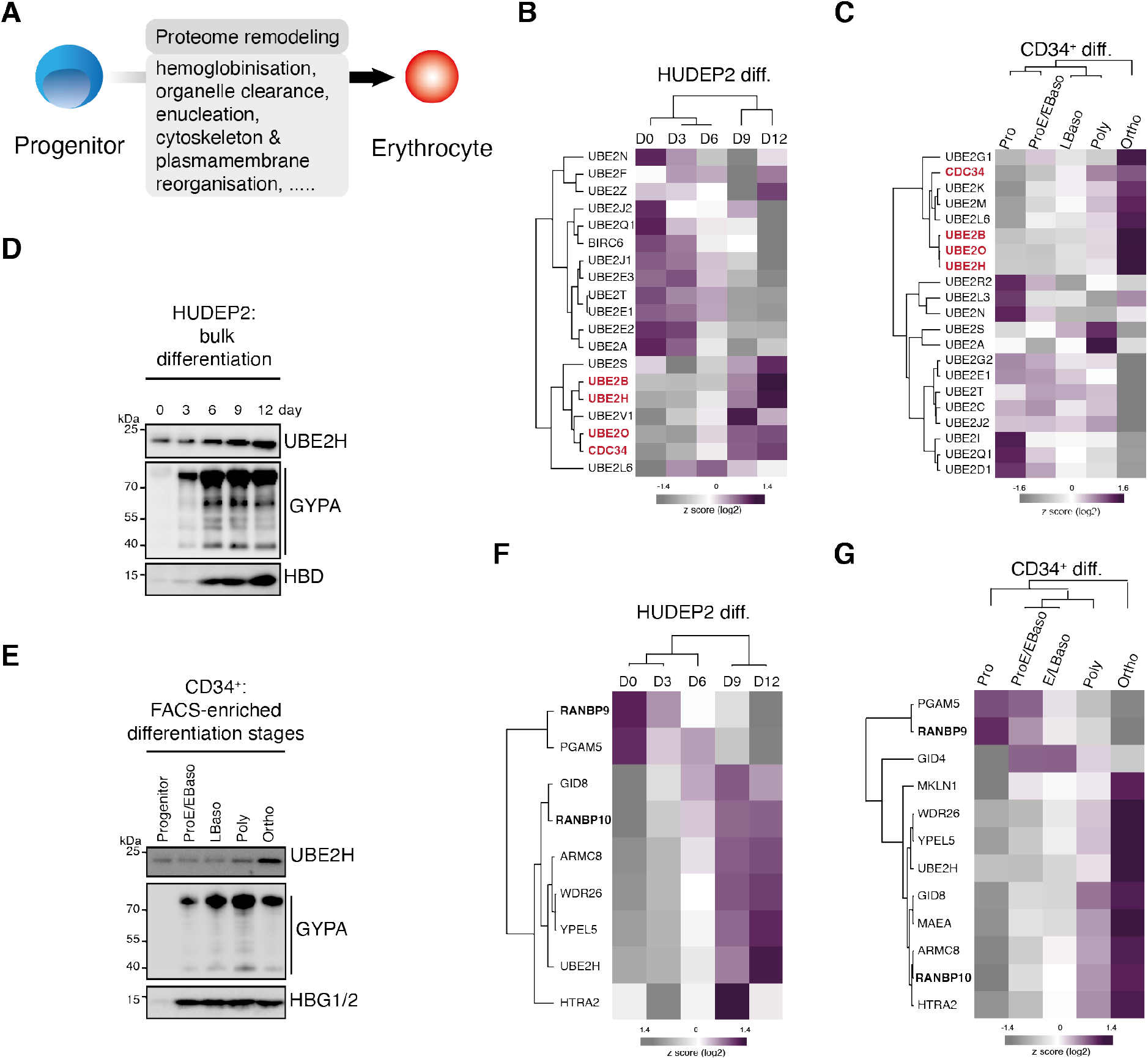
In-depth proteome profiling reveals stage-dependent expression of UBE2H and CTLH complex subunits in erythropoiesis. **A)** Cartoon indicating key features of mammalian erythropoiesis. **B)** Heat map of z-scored protein abundance (log2 DIA intensity) of differentially expressed E2 enzymes in differentiated HUDEP2 cells. **C)** Heat map of z-scored protein abundance (log2 DIA intensity) of differentially expressed E2 enzymes in differentiated CD34^+^ cells. **D)** HUDEP2 cells were differentiated *in vitro* and analysed by immunoblotting with indicated antibodies. **E)** CD34^+^ cells were differentiated *in vitro*, cell populations enriched by FACS and analysed by immunoblotting with indicated antibodies. **F)** Heat map of z-scored protein abundance (log2 DIA intensity) of differentially expressed CTLH complex subunits in differentiated HUDEP2 cells. **G)** Heat map of z-scored protein abundance (log2 DIA intensity) of differentially expressed CTLH complex subunits in differentiated CD34^+^ cells.

Our current knowledge of protein dynamics during erythropoiesis has been deduced largely from epigenetic and transcriptomic studies (reviewed in (An *et al*, 2015)), which used *in vitro* differentiation systems where erythroid progenitors, such as primary multipotent CD34^+^ haematopoietic stem and progenitor cells (HSPC) or immortalized CD34^+^-derived lines (known as HUDEP2 and BEL-A) (Kurita *et al*, 2013; Trakarnsanga *et al*, 2017), possess an autonomous differentiation programme with a capacity to complete terminal differentiation when cultured with cytokines and other factors (Seo *et al*, 2019). However, the dynamics of mRNA expression during erythropoiesis does not accurately predict protein expression (Gautier *et al*, 2016). We recently mapped the erythroid proteome landscape of defined precursors generated by *in vitro* differentiation of normal donor CD34^+^ cells (Karayel *et al*, 2020). Our findings provided insight into cellular remodelling at protein resolution and indicated high level of post-transcriptional regulation.

Ubiquitin (UB)-mediated protein degradation pathways are likely to play prominent roles in post-transcriptional regulation of erythropoiesis. Components of the E3 ubiquitin ligation machinery and deubiquitylases have been implicated in regulating protein stability and turnover in erythroid cell proliferation and maturation (Liang *et al*, 2019; Maetens *et al*, 2007; Mancias *et al*, 2015; Minella *et al*, 2008; Randle *et al*, 2015; Thom *et al*, 2014; Xu *et al*, 2020). The E2 enzyme UBE2O is greatly upregulated in reticulocytes and required for clearing ribosomes (Nguyen *et al.*, 2017). Recently, a functional role of the multi-protein *C*-*t*erminal to *L*is*H* (CTLH) E3 ligase complex was implicated in mammalian erythropoiesis. The CTLH subunits MAEA and WDR26 have been shown to be expressed in a differentiation stage-dependent manner and be implicated in maintaining erythroblastic islands in the bone marrow and regulating nuclear condensation in developing erythroblasts, respectively (Wei *et al*, 2019; Zhen *et al*, 2020). The tight correlation between protein abundance and functionality in differentiation suggested that large scale proteome profiling is a potential way to identify proteins that are important for the functional specialization of erythroid cells. Here, we profiled protein abundance of E2-E3 modules in erythroid differentiation uncovering a dynamic regulation of CTLH E3 ligase subunits and CTLH’s cognate E2 conjugating enzyme UBE2H. Interestingly, UBE2H amounts are dependent on active CTLH E3, suggesting a coupled E2/E3 regulation. We further show that CTLH complex compositions are remodelled and discrete complex assemblies are formed in a maturation stage-dependent manner. Our study indicates that unique UBE2H-CTLH assemblies are organized and co-regulated in functional E2-E3 modules and are required for the orderly progression of terminal erythroid maturation.

## RESULTS

### Stage-dependent expression of UBE2H and CTLH complex subunits in erythropoiesis

Reshaping of the erythropoietic proteome is thought to be regulated in part by the transient presence of stage-specific E2-E3 ubiquitin targeting machineries. To identify potential E2-E3 components, we applied a system-wide approach and established differentiation stage-specific proteomes of human erythropoiesis from two *in vitro* erythropoiesis cell model systems: CD34^+^ and HUDEP2 cells (Figure 1 **supplemental figure 1A 1B**). We recently described the stage-specific proteomes from *in vitro* differentiated CD34^+^ cells (Karayel *et al.*, 2020). HUDEP2 cells proliferate in immature progenitor state and can be induced to undergo terminal erythroid differentiation by modulating cell culture conditions (Figure 1 **supplemental figure 1B**) (Kurita *et al.*, 2013). In this study, HUDEP2 cells were shifted to differentiation conditions and semi-synchronous bulk cell populations were obtained at different time points (day 0, 3, 6, 9, and 12) corresponding to maturation stages spanning from proerythroblast to orthochromatic stages. Each population was processed in three biological replicates, and their proteomes were acquired by measuring single 100-minute gradient runs for each sample/replicate in data-independent acquisition (DIA) mode (Aebersold & Mann, 2016; Gillet *et al*, 2012; Karayel *et al.*, 2020; Ludwig *et al*, 2018). We measured HUDEP2 proteomes by DIA, as it provides sensitive and in-depth stage-specific proteome analysis of human erythropoiesis by maintaining a broad dynamic range of peptide detection in the presence of very abundant species, such as haemoglobin peptides, that accumulate to very high levels at late erythroid maturation stages (Karayel *et al.*, 2020). DIA raw files were searched with direct DIA (dDIA), yielding 6,727 unique proteins and quantitative reproducibility with Pearson correlation coefficients higher than 0.9 between the biological replicates of all populations (Figure 1 **supplemental figure 1C, 1D and supplemental table 1**). When we clustered the 2,771 differentially expressed proteins (ANOVA, FDR<0.01 and S0=0.1) we observed dynamic changes of the proteome between early (day 0) and late (day 12) time points across erythroid differentiation (Figure 1 **supplemental figure 1E**). The majority of proteins cluster into two co-expression profiles: continuous decrease or increase of protein levels that ultimately resulted in a reshaped erythrocyte-specific proteome.

We next examined our proteome data for all (~40) annotated human E2 conjugating enzymes. The levels of most E2s detected in HUDEP2 cells varied across maturation, with a cluster of six enzymes progressively accumulating until day 12 (Figure 1B). Included among them was UBE2O, which mediates ribosomal clearance in reticulocytes (Nguyen *et al.*, 2017). We expanded the analysis to stage-specific proteomes from *in vitro* differentiated CD34^+^ cells (Figure 1C) (Karayel *et al.*, 2020), which revealed a similar cluster of E2s upregulated at poly- and orthochromatic stages. Notably, UBE2B, UBE2O, CDC34 (aka UBE2R1), and UBE2H enzymes exhibited similar protein abundance profiles during HUDEP2 and CD34^+^ maturation suggesting important roles for these E2s during terminal erythropoiesis. CDC34, the cognate E2 for cullin-1 RING ligase (CRL1) complexes, is essential for cell cycle regulation (Kleiger *et al*, 2009; Skaar & Pagano, 2009). UBE2B (aka RAD6B) regulates DNA repair pathways, histone modifications, and proteasomal degradation (Kim *et al*, 2009; Varshavsky, 1996; Watanabe *et al*, 2004). We focused on UBE2H because it is transcriptionally regulated by the essential erythroid nuclear protein TAL1 and it accumulates to high levels during terminal maturation (Lausen *et al*, 2010; Wefes *et al*, 1995). Immunoblotting confirmed UBE2H protein upregulation during maturation of HUDEP2 and CD34^+^ cells, which was paralleling induction of the erythroid membrane protein CD235a (Glycophorin A, GYPA) and haemoglobin expression (Figure 1D **and** 1E).

The stage-dependent regulation of UBE2H suggested that a cognate E3 partnering with UBE2H would have a similar expression profile during erythropoiesis. *In vitro* ubiquitylation reactions indicate that UBE2H is the preferred E2 of the CTLH E3 complex (Lampert *et al*, 2018; Sherpa *et al*, 2021). The multi-protein CTLH complex consists of at least RANBP9 and/or RANBP10 (orthologous subunit of yeast Gid1), TWA1, ARMC8, WDR26 and/or MKLN1 (orthologous subunit of yeast Gid7), the catalytic module – MAEA and RMND5a that mediate ubiquitin transfer, and the substrate receptor GID4 (Kobayashi *et al*, 2007; Lampert *et al.*, 2018; Mohamed *et al*, 2021; Sherpa *et al.*, 2021; Umeda *et al*, 2003). Our analyses of the stage-dependent proteomes of differentiated HUDEP2 (Figure 1F) and CD34^+^ cells (Figure 1G) revealed that protein levels of most annotated CTLH subunits increased during erythroid maturation, in parallel with UBE2H. Interestingly, the homologues RANBP9 and RANBP10 showed an inverse expression pattern: RANBP9 levels were high at progenitor stages and dropped at later stages, whereas RANBP10 levels exhibited the opposite pattern. Taken together, analyses of differentiation-resolved proteomes revealed stage-dependent expression of CTLH subunits and UBE2H suggesting a dynamic assembly of distinct CTLH complexes linked to erythrocyte development.

### RANBP9 and RANBP10 assemble into distinct CTLH E3 complexes

To monitor CTLH complex assemblies, we next fractionated whole cell lysates from non-differentiated (day 0) or differentiated (day 6) HUDEP2 cells on 5-40% sucrose density gradients and detected CTLH subunits by immunoblot analysis. All CTLH subunits sedimented at ≥660 kDa, apparently corresponding to the supramolecular CTLH assemblies we previously described (Figure 2A) (Sherpa *et al.*, 2021). However, RANBP9 amounts in the CTLH fraction were higher at day 0 compared to day 6, while RANBP10 amounts had the opposite pattern, suggesting stage-specific modulation of CTLH complex composition and/or stoichiometry during differentiation. To further test this, we immunoprecipitated (IP) CTLH complexes from day 0 and day 6 lysates using an anti-ARMC8-specific nanobody. This resulted in coprecipitation of predominantly RANBP9 from day 0 cell lysates, whereas RANBP10 was found in elevated levels in lysates from day 6 differentiated HUDEP2 cells (Figure 2B). Next, we asked whether RANBP9 and RANBP10 can independently form CTLH complexes. Using CRISPR-Cas9 editing, we deleted either *RANBP9* or *RANBP10* in HUDEP2 cells (Figure 2C). Notably, we observed elevated RANBP10 amounts in *RANBP9^-/-^* cells, and slightly increased RANBP9 amounts in *RANBP10^-/-^* cells, suggesting a possible reciprocal compensatory effect for the loss of either homologue. Whole cell lysates of parental and KO lines were analysed using sucrose density gradients and immunoblot analysis to assess the sedimentation of RANBP9 and RANBP10 (Figure 2D). The sedimentation of the supramolecular CTLH complex containing RANBP9 was similar in parental and *RANBP10^-/-^* cells. Likewise, RANBP10-CTLH assemblies sedimented similarly in parental and *RANBP9^-/-^* cells. These data indicate that RANBP9 and RANBP10 assemble in distinct CTLH E3 supramolecular complexes.

**Figure 2.**
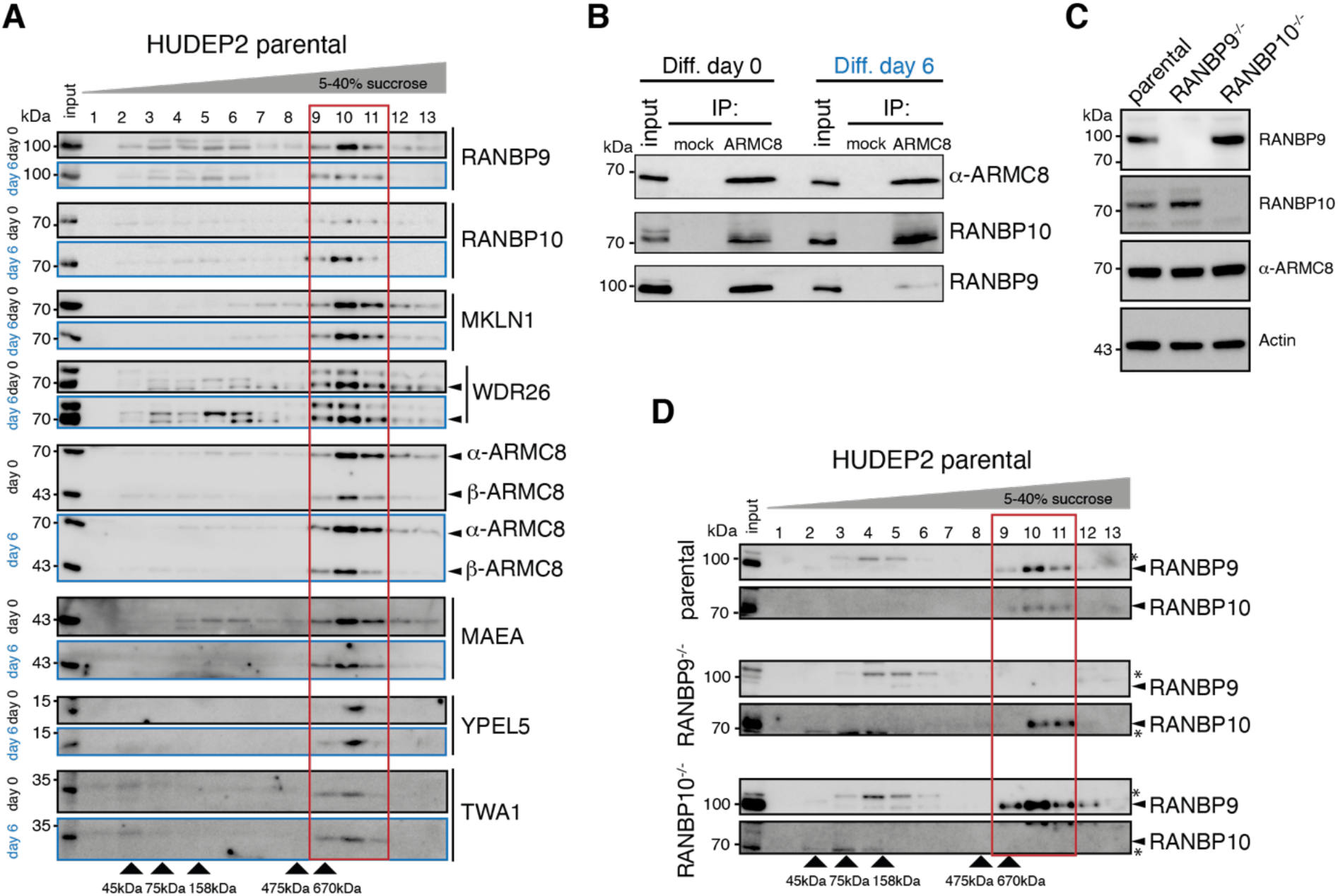
RANBP9 and RANBP10 assemble in distinct CTLH E3 complexes. **A)** HUDEP2 cell lysates from differentiation day 0 and day 6 were separated on sucrose gradients, and fractions analysed by immunoblotting with indicated antibodies. **B)** Immunoprecipitation (IP) of HUDEP2 cell lysates from differentiation day 0 and day 6 with IgG control (mock) and ARMC8-specific nanobody, and immunoblot analysis with indicated antibodies. **C)** Immunoblots of lysates of HUDEP2 parental, RANBP9^-/-^, and RANBP10^-/-^ cells probing for RANBP9 and RANBP10. Actin serves as loading control. **D)** Sucrose gradient fractionation of HUDEP2 cell lysates from RANBP9^-/-^ or RANBP10^-/-^ knock out lines, fractions were analysed by immunoblotting with indicated antibodies.

### RANBP9 and RANBP10 form homologous CTLH E3 complexes that cooperate with UBE2H to promote ubiquitin transfer

Recent cryo-EM maps of human CTLH sub- and supramolecular complexes revealed that RANBP9 supports complex assembly (Sherpa *et al.*, 2021). Beyond an N terminal extension unique to RANBP9, RANBP9 and RANBP10 have a common domain architecture (Figure 3 **supplemental figure 2A**). Hence, we reasoned that RANBP10 may form similar parallel interactions with other CTLH subunits. To test this, we expressed and purified a core CTLH subcomplex, containing a scaffold module (RANBP10, TWA1, α-ARMC8), the catalytic module (MAEA and RMND5a), and the substrate receptor GID4 (named thereafter RANBP10-CTLH^SR4^). In parallel, we generated the previously described homologous complex where RANBP9 replaced RANBP10 (RANBP9-CTLH^SR4^) (Mohamed *et al.*, 2021; Sherpa *et al.*, 2021). The two complexes eluted at similar range in size exclusion chromatography (SEC), indicating they had comparable subunit stoichiometry (Figure 3A). Cryo-EM analysis of the RANBP10-CTLH^SR4^ peak fraction yielded a reconstitution at ~12 Å resolution (Figure 3B). Comparison to the previously determined RANBP9-CTLH^SR4^ map (EMDB: EMD-12537) revealed an overall structural similarity and the clamp-like assembly of substrate receptor scaffolding (SRS) and catalytic (Cat) modules conserved in related yeast GID complexes (Qiao *et al*, 2020; Sherpa *et al.*, 2021) (Figure 3 **supplemental figure 1**). Furthermore, atomic coordinates of α-ARMC8, hGID4 and TWA1 derived from the RANBP9-CTLH^SR4^ structure (PDB: 7NSC), along with crystal structure of RANBP10-SPRY domain (PDB: 5JIA) fit into the 7.3 Å resolution focused refined map of RANBP10-CTLH^SR4^ map (Figure 3C). To accurately position RANBP10, the crystal structure of the RANBP10’s SPRY domain (PDB: 5JIA) was superimposed to the structure of the recently published RANBP9-CTLH^SR4^ (Figure 3 **supplemental figure 2A and B**). Thus, at an overall level, the RANBP10-CTLH^SR4^ and RANBP9-CTLH^SR4^ complexes are structurally homologous.

**Figure 3.**
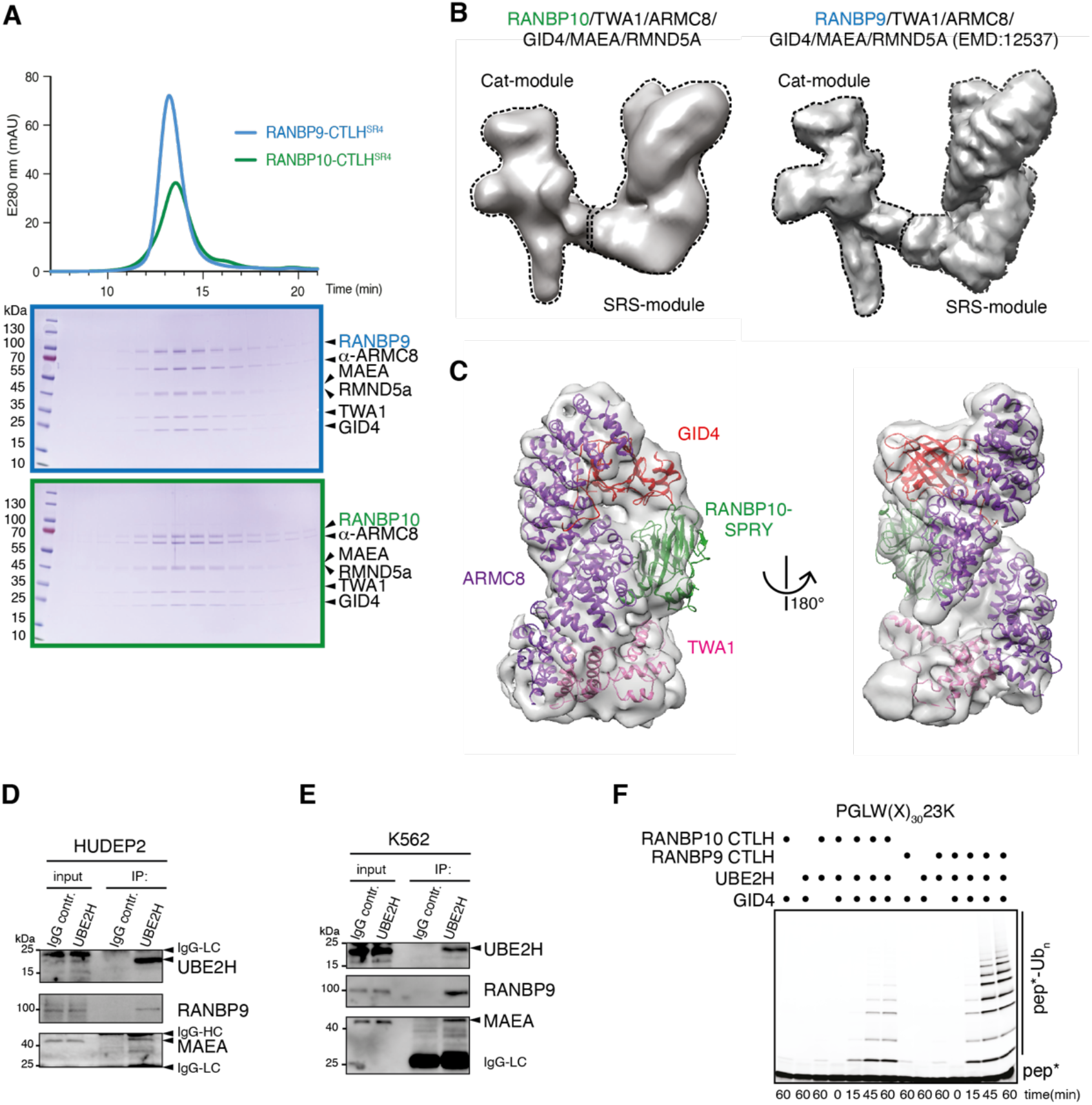
RANBP9 and RANBP10 form homologous CTLH E3 complexes that promote ubiquitin transfer in cooperation with UBE2H. **A)** Chromatograms (top) and Coomassie-stained SDS PAGE gels (bottom) from size exclusion chromatography of recombinant RANBP10-CTLH^SR4^ and RANBP9-CTLH^SR4^ complexes. **B)** Cryo-EM map of RANBP10-CTLH^SR4^ (left) and RANBP9-CTLH^SR4^ (EMD:12537) (right) with Cat-module and SRS-module indicated. **C)** Focused refined map of the RANBP10-CTLH SRS-module with coloured subunits: ARMC8, purple; TWA1, salmon; hGid4, red; RANBP10 SPRY-domain, green; **D)** Immunoprecipitation (IP) from HUDEP2 cell lysates with IgG control and UBE2H-specific antibody and immunoblot analysis. **E)** Immunoprecipitation (IP) from K562 cell lysates with IgG control and UBE2H-specific antibody and immunoblot analysis. IgG light chain (IgG-LC), IgG heavy chain (IgG-HC). **F)** Fluorescence scan of SDS-PAGE gels presenting time course of *in vitro* ubiquitylation assay with fluorescently-labelled model substrate peptide PGLW(X)_n_-23K with lysine at position 23 (pep*), in the presence of UBE2H, RANBP10-CTLH or RANBP9-CTLH, and GID4.

We next asked whether the structural similarity of RANBP10-CTLH^SR4^ and RANBP9-CTLH^SR4^ complexes extended to the mechanism of ubiquitin transfer activity. First, we assessed the physical association between UBE2H and CTLH complex subunits in HUDEP2 or erythroleukemia K562 cells using anti-UBE2H IPs (Figure 3D 3E) (Andersson *et al*, 1979). Endogenous UBE2H specifically co-precipitated the Cat-module subunit MAEA in whole cell lysates from both cell lines, indicating that UBE2H can form a reasonably stable E2-E3 enzyme module. Next, we tested ubiquitin transfer activity by *in vitro* ubiquitylation assays with a fluorescently labelled model peptide substrate (Sherpa *et al.*, 2021). This model substrate consisted of an N-terminal PGLW sequence, that binds human GID4 (Dong *et al*, 2020; Dong *et al*, 2018), and a 30 residue linker sequence with the target lysine (K) towards the C-terminus at position 23 (PGLW[X]_30_K23) (Figure 3F). In a reaction with UBE2H, both RANBP9-CTLH and RANBP10-CTLH promoted polyubiquitylation of the model substrate peptide in a GID4-dependent manner, although RANBP10-CTLH was less active under these conditions than the homologous RANBP9-CTLH complex. Cumulatively, biochemical and structural data revealed that RANBP9 and RANBP10 can assemble in distinct homologous CTLH complexes capable of activating UBE2H-dependent ubiquitin transfer activity. More broadly, CTLH may represent a larger family of E3 ligase complexes generated by assembly of different variable members with invariable core scaffold subunits.

### Catalytically inactive CTLH E3 complexes and UBE2H deficiency cause deregulated proteome dynamics

To investigate potential functional roles of CTLH E3 complex assemblies and UBE2H in erythroid cells, we edited HUDEP2 cell lines with CRISPR-Cas9 to disrupt either *UBE2H* or *MAEA* (Figure 4 **supplemental figure 1A and 1B**). Presumably, deletion of *MAEA*, the enzymatic subunit in all CTLH E3 assemblies, will result in a complete loss of all CTLH complex E3 ligase activities. As terminal erythropoiesis is characterized by a stage-dependent proteome remodelling (Figure 1 **supplemental figure 1**)(Gautier *et al.*, 2016; Karayel *et al.*, 2020), we first assessed how UBE2H and MAEA deficiency might alter the global proteome. Global proteomic analysis of undifferentiated parental, *UBE2H^-/-^* (clone 13), and *MAEA^-/-^* (clone 3-1) HUDEP2 cells identified 6,210 unique proteins in total (Figure 1 **supplemental figure 1C and 1D and supplemental table 2**) (Student t-test with FDR<0.05 and S0=0.1). We found that 18% (1,170) and 6.5% (404) of all proteins were significantly changed in *UBE2H^-/-^* vs. parental and *MAEA^-/-^* vs. parental comparisons, respectively (Students t-test with FDR<0.05 and S0=0.1)(Figure 4A and 4B). Notably, 271 of these proteins were differentially changed in both, a MAEA- and UBE2H-dependent manner (Figure 4C). These included several erythroid-specific proteins including haemoglobin subunits (HBD, HBG2, and HBM) and Band3 (SLC4A1) in both comparisons indicating an erythroid-typical remodelled proteome (Figure 4D). In Gene Ontology (GO) enrichment analysis proteins associated with annotations related to erythropoiesis such as ‘haemoglobin complex’ and ‘oxygen binding’ were significantly enriched in both *MAEA^-/-^* and *UBE2H^-/-^* cells compared to parental cells (Figure 4E). Next, we turned to network analysis of physically interacting or functionally associated proteins that were significantly up-regulated in both *MAEA^-/-^* and *UBE2H^-/-^* cells compared to parental. Overrepresentation analysis revealed significant enrichment of protein networks involved in terms including ‘haemoglobin-haptoglobin complex’ and ‘regulation of erythrocyte differentiation’ (Benjamini Hochberg, FDR 5%) (Figure 5F). Consistent with accelerated erythroid maturation, we detected a 3-6-fold higher levels of cellular heme compared to parental cells in lysates of *MAEA^-/-^* and *UBE2H^-/-^* cells (Figure 4G). To test the temporal nature of the proteome changes, we obtained and compared differentiation time-specific proteomes (day 0 to day 12) of parental and *MAEA^-/-^* (clone 3-1) cells (Figure 1 **supplemental figure 1C and** Figure 4 **supplemental figure 2A and 2B**). The five distinct temporal stages of erythroid differentiation clustered separately by principal component analysis (PCA) with high consistencies between the three biological replicates (Figure 4H). Remarkably, parental versus *MAEA^-/-^* clusters progressively diverged up to differentiation day 6 and remained separated to day 12, suggesting a MAEA-dependent proteome remodelling at early stages of differentiation (Figure 5H). PCA analysis based on 28 erythroid-specific marker proteins (Figure 4I) revealed that the separation of parental versus *MAEA^-/-^* cluster was pronounced even at early differentiation time points day 0 (Figure 4J). In fact, the *MAEA^-/-^* cluster at day 0 shows a closer correlation with the parental cluster at day 3 than day 0. Together, *UBE2H^-/-^* and *MAEA^-/-^* cells are characterized by enhanced proteome-wide changes towards an erythroid-specific proteome signature.

**Figure 4.**
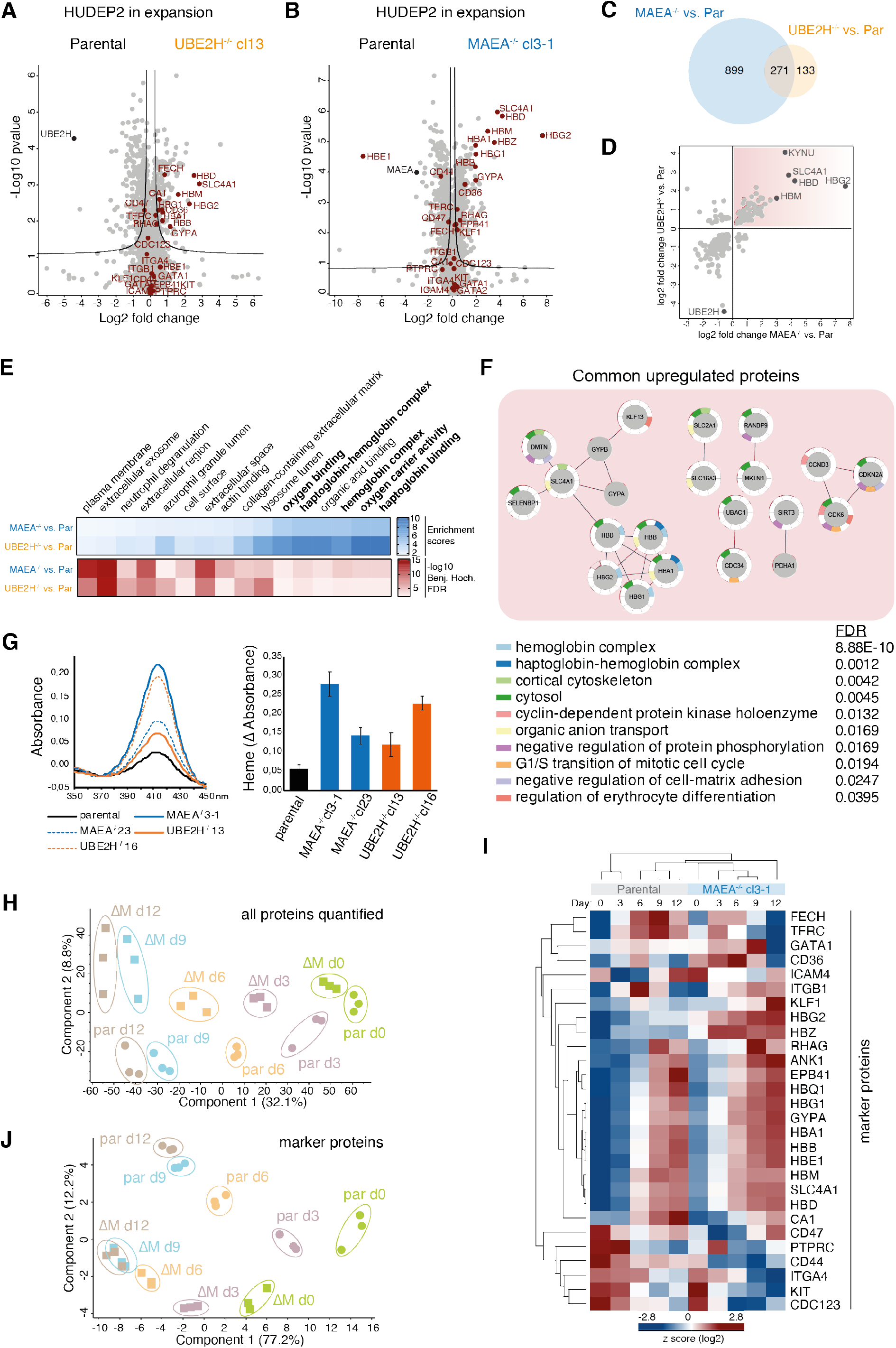
Catalytically inactive CTLH E3 complexes and UBE2H deficiency cause deregulated proteome dynamics. **A)** Volcano plots of p-values (-log10) versus protein abundance (log2) differences between parental and UBE2H^-/-^cl13 cells with erythroid marker proteins highlighted in red. **B)** Volcano plots of p-values (-log10) versus protein abundance (log2) differences between parental and MAEA^-/-^cl3-1 cells with erythroid marker proteins highlighted in red. **C)** Overlap between proteins with abundance differences of UBE2H^-/-^ versus parental and MAEA^-/-^ versus parental comparisons. **D)** Proteins with abundance differences (log2) of UBE2H^-/-^ versus parental blotted against proteins with abundance differences (log2) MAEA^-/-^ versus parental comparisons. Commonly enriched proteins are highlighted (top right quadrant). **E)** Gene Ontology (GO) enrichment analyses of up-regulated protein in MAEA^-/-^ versus parental and UBE2H^-/-^ versus parental comparisons performed using Fisher’s exact test (Benjamini-Hochberg, FDR 5%). **F)** Overrepresentation analysis revealed the significant enrichment of protein networks (Benjamini-Hochberg, FDR 5%) based on physically interacting or functionally associated proteins which were significantly up-regulated in both MAEA^-/-^ and UBE2H^-/-^ versus parental cells. **G)** Spectra of cleared cell lysates of indicated cell lines showing Soret absorbance peak at 414 nm corresponding to heme-bound haemoglobins (left). Relative Soret absorbance peak intensity calculated from spectra of cleared cell lysates of indicated cell lines. Results are mean ± SD of n=3 experiments. **H)** Principal Component Analysis (PCA) of erythroid differentiation stages (day 0, green; day 3, purple; day 6, orange; day 9, blue; day 12, brown) of HUDEP2 parental (par) and MAEA^-/-^cl3-1 (∆M) cell lines with their biological replicates based on expression profiles of all quantified proteins. **I)** Heat map of z-scored protein abundance (log2 DIA intensity) of differentially expressed erythroid marker proteins in differentiating HUDEP2 parental and MAEA^-/-^cl3-1 cells. **J)** PCA as in H), but based on expression profiles of selected erythroid marker proteins in I).

**Figure 5.**
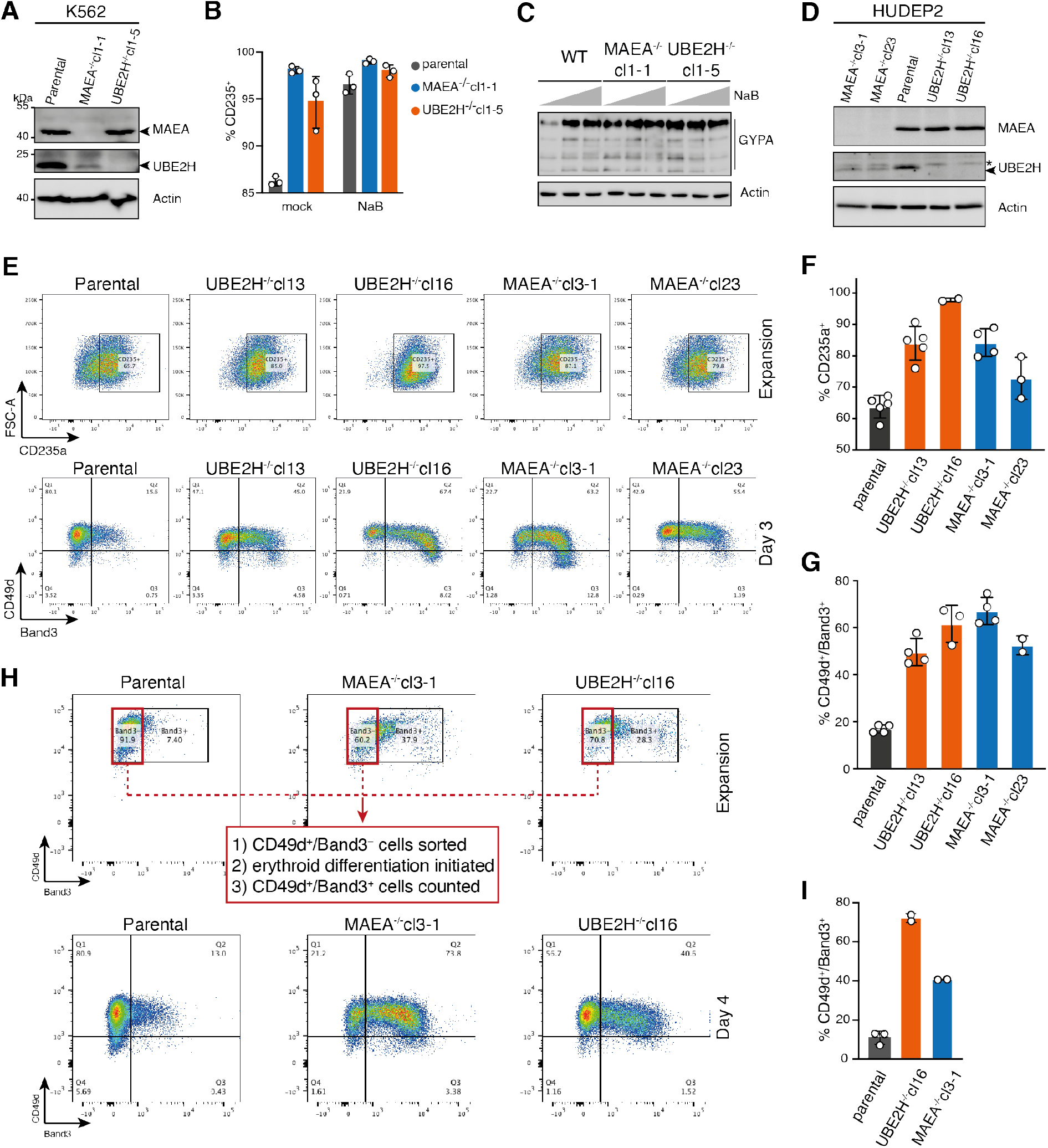
MAEA and UBE2H deficiency cause aberrant erythroid differentiation. **A)** Immunoblots of lysates of K562 parental, MAEA^-/-^cl1-1, UBE2H^-/-^cl1-5 cells probing for MAEA and UBE2H. Actin serves as loading control. **B)** Graph shows fraction of CD235a-expressing K562 parental, MAEA^-/-^cl1-1, UBE2H^-/-^cl1-5 cells in the presence of Na-butyrate (NaB) or mock treated. Error bars represent mean ±STDEV of n=3 biological replicates. **C)** Immunoblot analysis of cell lysates from K562 parental, MAEA^-/-^cl1-1, UBE2H^-/-^cl1-5 treated with 0, 0.3, or 0.6 mM NaB for 24 hrs. Anti-GYPA/CD235a antibody detects variant forms of glycosylated GYPA. Actin serves as loading control. **D)** Immunoblots of lysates of HUDEP2 parental, MAEA^-/-^, and UBE2H^-/-^ knock out clones probing for MAEA and UBE2H. Actin serves as loading control. **E)** Flow cytometry blots of indicated HUDEP2 cell lines showing CD235a (top) and CD49d/Band3 (bottom) expression in expansion growing media condition(top) or day 3 after induced erythroid differentiation (bottom). **F)** Quantitation of **E)** with graph showing fraction of CD235a^+^ cells. Error bars represent mean ± SD of n=2 to 5 biological replicates. G) Quantitation of E) with graph showing fraction of CD49d^+^/Band3^+^ cells. Error bars represent mean ± SD of n=2 to 4 biological replicates. H) Indicated HUDEP2 cell lines, cultured in expansion media, were sorted for CD49d^+^/Band3^-^ (flow cytometry blots, top) followed by induction of erythroid maturation. Flow cytometry blots of indicated HUDEP2 cell lines showing CD49d/Band3 expression at day 4 after induced erythroid maturation. I) Quantitation of H) with graph showing fraction of CD49d^+^/Band3^+^ cells. Error bars represent mean ± SD of n=2 biological replicates.

### MAEA and UBE2H deficiency cause aberrant erythroid differentiation

The deregulated proteome landscapes of *UBE2H^-/-^* and *MAEA^-/-^* cells indicate a functional role of UBE2H-CTLH modules in the initiation of erythroid differentiation of progenitors and/or the progression through erythropoiesis. First, we used K562 cells as a surrogate erythroid cell model that expresses erythroid markers, including CD235a/GYPA and haemoglobins upon treatment with the pan histone deacetylase inhibitor Na-butyrate (NaB) (Andersson *et al.*, 1979). We generated K562 *UBE2H^-/-^* and *MAEA^-/-^* cells by CRISPR-Cas9 editing (Figure 5A and Figure 4 **supplemental figure 1B**). Parental and knock out cell lines were either mock or NaB-treated and erythroid differentiation was assessed by CD235a/GYPA surface expression via flow cytometry. At baseline cell conditions (mock) *MAEA*^-/-^ and *UBE2H*^-/-^ lines showed increased CD235a/GYPA positive (CD235a^+^) cells comparable to NaB-treated parental cells (Figure 5B and Figure 5 **supplemental figure 1C**). Furthermore, immunoblot analysis showed increased CD235a/GYPA expression in whole cell lysates of *MAEA*^-/-^ and *UBE2H*^-/-^ lines treated at low dose of NaB suggesting that *MAEA* and *UBE2H* deficiency might promote erythroid differentiation (Figure 5C). To further substantiate the observation, we expanded the analysis to *MAEA-* and *UBE2H*-deficient HUDEP2 cell lines. Flow cytometry analysis revealed an elevated proportion of CD235^+^ cells in clones lacking *MAEA* (cl3-1 and cl23) or *UBE2H* (cl13 and cl16), indicating spontaneous erythroid maturation in expansion medium (Figure 5E and 5F). Next, we induced erythroid differentiation for three days and evaluated erythroid markers CD49d (Integrin alpha 4) and Band3 (SLC4A1). Each *MAEA^-/-^* and *UBE2H^-/-^* clone showed higher CD49d^+^/Band3^+^ cell populations compared to parental HUDEP2, indicating either precocious or accelerated maturation (Figure 5E and 5G). To determine whether *MAEA^-^/-* and *UBE2H^-/-^* cells mature faster compared to controls, we sorted Band3^-^ cells to generate “synchronous” non-differentiated populations prior to differentiation. CD49d^+^/Band3^+^ measurement after day 4 confirmed an accelerated maturation in the absence of UBE2H or MAEA (Figure 5H and 5I). Taken together, the data suggest that UBE2H and MAEA deficiencies share the common defects of spontaneous erythroid differentiation of cells while propagated in expansion media and accelerated initial stages of erythroid maturation.

### Cellular abundance of UBE2H is coupled to functional MAEA

Evidence for a regulatory relationship between UBE2H and MAEA during erythropoiesis was provided by an unexpected observation that UBE2H protein levels are dependent on MAEA. Proteomics and immunoblot analyses showed consistently lower UBE2H levels in K562 and HUDEP2 *MAEA^-/-^* cell lines (Figure 4D and Figure 5A and 5D). Moreover, differentiating *MAEA^-/-^* cells failed to express increased UBE2H protein levels at terminal maturation stages (day 9 and 12) (Figure 6A). UBE2H mRNA levels, however, were not significantly different between parental and *MAEA^-/-^* cells at day 9 of differentiation (Figure 6B), indicating that transcriptional regulation of UBE2H is not affected. We next expanded our analysis to the K562 cells (Figure 6C). Parental and knock-out K562 cell lines were ether treated with NaB to induce erythroid-like or with 12-O-Tetradodecanoyl-phorbol-13 acetate (TPA, chemical activator of PKC kinase) to induce megakaryocyte-like differentiation (Tabilio *et al*, 1983). Both, NaB and TPA efficiently induced UBE2H protein levels in parental cells suggesting that UBE2H regulation is not restricted to erythroid differentiation (Figure 6D and 6E). In contrast, *MAEA^-^/-* cells had constitutively less and only marginally induced UBE2H protein levels in response to NaB or TPA. Ectopic expression of Myc-tagged MAEA in *MAEA^-/-^* cells rescued UBE2H protein levels, confirming that the presence of MAEA is critical for maintaining UBE2H protein levels (Figure 6 **supplemental figure 1A**). Notably, cells either lacking CTLH’s substrate receptor GID4 or subunits of the supramolecular module, WDR26 and MKLN1, showed UBE2H abundance and regulation similar to parental cells (Figure 6D, 6E, and Figure 6 **supplemental figure 1D**). The assessment of UBE2H mRNA levels revealed no significant difference of NaB-induced UBE2H transcription between parental and MAEA^-/-^ cells (Figure 6F).

**Figure 6.**
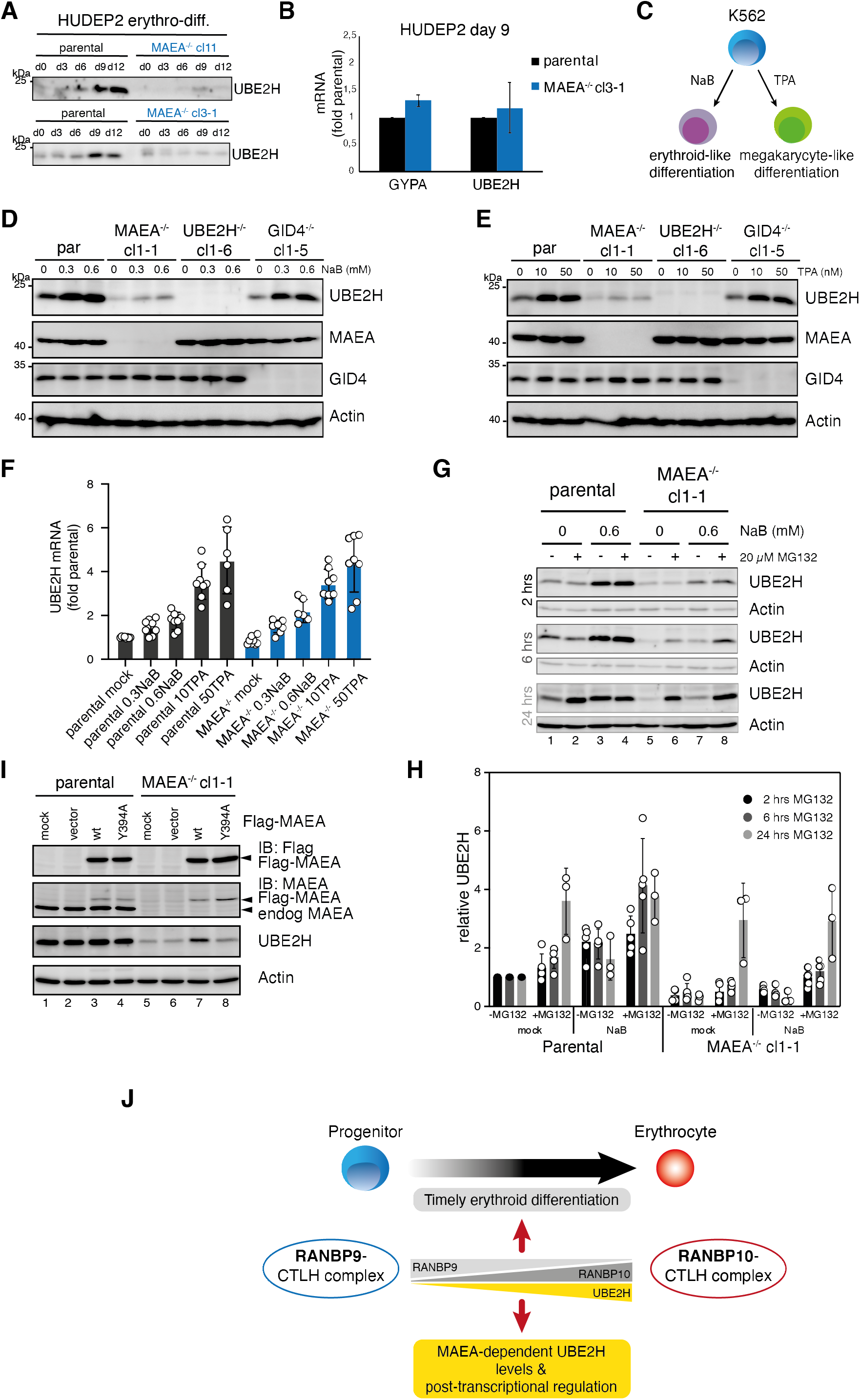
Cellular abundance of UBE2H is coupled to functional MAEA. **A)** HUDEP2 parental and MAEA^-/-^ (clone 3-1 and 11) cells were differentiated *in vitro* and analysed by immunoblotting to detect UBE2H protein levels. **B)** mRNA determination by RT-qPCR of HUDEP2 parental and MAEA^-/-^cl3-1 cells at differentiation stage day 9. Results (normalized to GAPDH) are mean ± SD of n=2 experiments. **C)** K562 cell can be either induced with Na-butyrate (NaB) for erythroid-like differentiation, or induced with 12-O-Tetradodecanoyl-phorbol-13 acetate (TPA) for megakaryocyte-like differentiation. **D)** K562 parental and knock out cell lines were treated with NaB for 24 hrs and analysed by immunoblotting with indicated antibodies. **E)** K562 parental and knock out cell lines were treated with TPA for 24 hrs and analysed by immunoblotting with indicated antibodies. **F)** K562 parental and MAEA^-/-^cl1-1 cell lines were treated with either NaB or TPA for 24 hrs, and UBE2H mRNA levels determination by RT-qPCR. Results (normalized to GAPDH) are mean ± SD of n=6 to 8 experiments. G) K562 parental and MAEA^-/-^cl1-1 cell lines were mock or 0.6 mM NaB treated for 24 hrs, followed by 2, 6, and 24 hours proteasome inhibition with 20 μM MG132 and immunoblot analysis of cell lysates for UBE2H. Actin serves as protein loading control. H) Quantitation of UBE2H immunoblot signals from G) normalized to Actin and relative to parental mock values. Graph shows results by mean ± SD of n=3 to 5 experiments. I) K562 parental and MAEA^-/-^cl1-1 cells were mock, empty vector, Flag-MAEA wildtype (WT), or mutated Flag-MAEA-Y394A (Y394A) transfected and cell lysates were analysed for UBE2H protein levels by immunoblotting. J) Model indicating stage-dependent CTLH complex assemblies coupled with UBE2H abundance required in erythropoiesis.

To further investigate the mechanism that underlies regulation of UBE2H protein level and stability we first asked whether UBE2H is targeted by proteasomal degradation. K562 parental and *MAEA^-/-^* cells were mock or NaB-treated followed by a time course treatment with proteasomal inhibitor (MG132). Whereas MG132 treatment for 2 hours had only a modest effect on UBE2H levels, 6-24 hours exposure resulted in an increasing stabilisation of UBE2H in *MAEA^-/-^* cells (lane 6 and 8) matching levels of parental cells (lane 2 and 4) (Figure 6G and 6H). Notably, NaB-induced UBE2H levels are comparable to 24 hours MG132 treated parental cells. Second, we asked whether MAEA activity is required for UBE2H abundance and stability. To this end, we studied the MAEA Y394A mutation (MAEA-Y394A) that abolishes activity of the Cat-module *in vitro* (Sherpa *et al.*, 2021), but maintains binding capacity to UBE2H in immune precipitation (IP) experiments (Figure 6 **supplemental figure 1B and 1C**). Ectopic expression of Flag-tagged MAEA-Y394A in *MAEA^-/-^* cells revealed no rescued UBE2H protein levels, whereas wild type MAEA expression resulted in a significant increase of UBE2H abundance (compare lane 7 and 8) (Figure 6I). Hence, association with MAEA is itself not sufficient to maintain UBE2H protein levels, but rather the catalytic activity of MAEA is required. Overall, we conclude that UBE2H protein levels are coupled to the presence of active MAEA and are controlled by differentiation-induced transcription and post-transcriptionally by proteasomal degradation.

## DISCUSSION

We demonstrate here that in-depth analysis of dynamic proteome profiles obtained from different *in vitro* reconstituted erythropoiesis systems is an effective method to uncover differentiation stage-dependent expression of protein and protein complexes with functional roles in erythropoiesis. We observed that UBE2H and CTLH E3 complex assemblies form co-regulated E2-E3 modules required for erythropoiesis. Importantly, distinct protein profiles of the CTLH subunit homologues RANBP9 and RANBP10, suggest a remodelling of CTLH complex and the presence of stage-dependent CTLH assemblies.

The activity and substrate specificity of multi-subunit E3 ligases are generally regulated by the control of complex subunit assembly. One of the best studied E3 ligase complexes are members of the CRL family which engage interchangeable substrate receptor/adapter complexes for substrate-specific ubiquitylation. Substrate receptor assembly is kept in a highly dynamic state (Reitsma *et al*, 2017; Straube *et al*, 2017), whereby specific expression of substrate receptors in response to external and internal cellular cues, as well as in a cell-type and tissue specific way, allows the formation of CRLs for selective and efficient client substrate ubiquitylation (Gupta *et al*, 2013; McGourty *et al*, 2016; Ravenscroft *et al*, 2013). Recently, the activity of the multiprotein yeast GID (orthologue of human CTLH complex) was shown to be predominantly regulated by engaging interchangeable substrate receptors, which conceivably target distinct substrates for degradation (Chrustowicz *et al*, 2021; Kong *et al*, 2021; Liu & Pfirrmann, 2019; Melnykov *et al*, 2019; Shin *et al*, 2021)(Langlois et al. 2021 BioRxiv doi: https://doi.org/10.1101/2021.09.02.458684). However, the regulation of human CTLH activity via substrate receptor assembly is not well understood. Here we show that protein levels of GID4 – the only substrate receptor identified in higher eukaryotes – did not significantly change in the HUDEP2 differentiation system, suggesting that GID4 may not be the critical regulatory subunit of CTLH complex during erythropoiesis. In contrast, other CTLH subunits were significantly up or down regulated with correlating protein intensity profiles indicating distinct classes of CTLH assemblies. In particular, the homologues scaffold module subunits RANBP9 and RANBP10 displayed inverse expression profiles. As a consequence, non-differentiated HUDEP2 cells preferentially assembled RANBP9-CTLH complexes, whereas differentiated cells showed increased levels of RANBP10-CTLH complexes. These complexes were formed independently in cells, and additional structural/biochemical characterization of recombinant complexes revealed overall similar architectures of catalytic and scaffold modules exerting E3 ligase activities *in vitro* with UBE2H.

Despite the compelling evidence of distinct CTLH assemblies in differentiating erythroid cells, we can only speculate about the mechanism of assembly and remodelling of complex subunits. RANBP9 and RANBP10 are unlikely to exchange freely within the CTLH complexes given that both subunits form extended surface interactions with ARMC8 and TWA1 forming the core of the scaffold module (Sherpa *et al.*, 2021). Indeed, depleting RANBP9 destabilized CTLH in cell lysates (Maitland *et al*, 2019). Hence, RANBP9 and RANBP10 are likely to assemble CTLH complexes *de novo,* dependent on their availability and expression profiles. Furthermore, we observed increased protein levels of RANBP10 in *RANBP9^-/-^* cells (and increased RANBP9 in *RANBP10^-/-^* cells), suggesting that the cellular stoichiometry of CTLH subunits allows a “compensatory” assembly of RANBP10-CTLH complexes. Importantly, our description of RANBP9-CTLH and RANBP10-CTLH complexes substantiate the notion that CTLH complexes exist in multiple compositions and architectures thereby expanding the complexity of the CTLH E3 family (Sherpa *et al.*, 2021).

To determine the biological role of all possible CTLH complexes in erythropoiesis, we focused on *MAEA* knock-out cells. MAEA deficiency in K562 and HUDEP2 cells caused significantly lower protein amounts of RMND5a and UBE2H, suggesting the ubiquitin transfer activity of most - if not all - cognate UBE2H-CTLH modules was eliminated. Analysis of *MAEA* and *UBE2H* in HUDEP2 cells, that were cultured under expansion growing (non-differentiating) conditions, showed increased haemoglobinization and enrichment of erythroid marker proteins, resembling a spontaneously differentiated cell population. Hence, these knock out cells might be more susceptible to signals that promote erythropoiesis. Therefore, we reasoned that MAEA and UBE2H are required to maintain HUDEP2 cells in a dormant/quiescent progenitor stage. The *in vitro* reconstitution of erythropoiesis by the HUDEP2 system does not fully recapitulate *in vivo* erythropoiesis. However, our findings are supported by *MAEA* knock out studies in mice. Conditional *MAEA* deletion in murine haematopoietic stem cells (HSCs) impaired HSC quiescence, leading to a lethal myeloproliferative syndrome (Wei *et al*, 2021). The authors proposed a mechanism whereby the absence of MAEA leads to a stabilisation of several haematopoietic cytokine receptors causing prolonged intracellular signalling (Wei *et al.*, 2021). A similar concept might apply to the observed phenotypes of HUDEP2 *MAEA^-/-^* cells. Besides overrepresentation of several erythroid plasma membrane proteins, these cells have increased TFRC (Transferrin receptor 1) levels, and hence, are potentially more responsive to extracellular ferritin, enabling increased heme and haemoglobin production. Furthermore, mouse studies, that either conditionally deleted *MAEA* in central macrophages of erythroblastic islands or in erythroid progenitors, have revealed abnormal erythroblast maturation in the bone marrow showing altered profiles with distinct accumulation of maturation stages (Wei *et al.*, 2019). This phenotype is, in part, recapitulated in *MAEA-* and *UBE2H-*deficient HUDEP2 cells by an apparent accumulation of early maturation stages.

The functions of E2-E3 ubiquitylation modules are typically considered to be regulated via the E3 enzyme. However, E3 ubiquitylation involves different E2s which themselves can be regulated by multiple mechanisms (Stewart *et al*, 2016), such as modulation by transcriptional/translational control (Mejia-Garcia *et al*, 2015; Whitcomb *et al*, 2009; Ying *et al*, 2013). Apart from transcriptionally regulation of UBE2H in mammalian erythropoiesis (Lausen *et al.*, 2010; Wefes *et al.*, 1995), its *Drosophila* orthologue Marie Kondo (Kdo) was shown to be translationally upregulated upon oocyte-to-embryo transition (Zavortink *et al*, 2020), suggesting that UBE2H levels are regulatory nodes in developmental processes of higher eukaryotes. In agreement with transcriptional upregulation of UBE2H mRNA in terminal erythroid maturation (Lausen *et al.*, 2010), we observe a substantial increase in UBE2H protein at orthochromatic stages of differentiating HUDEP2 cells. Surprisingly, absence of MAEA caused reduced protein but not mRNA levels of UBE2H, suggesting a MAEA-dependent post-transcriptional mode of regulation. A MAEA mutant that still can bind UBE2H, but is defective in E3 ligase activity, does not efficiently rescue UBE2H levels in *MAEA^-/-^* cells. Therefore, in the absence of an active CTLH complex, UBE2H might become an “orphan” E2, which is subsequently targeted for proteasomal degradation. To our knowledge, UBE2H-CTLH is the first E2-E3 module described whereby E2 levels are coupled to the activity of the cognate E3.

Cumulatively, our work features a mechanism of developmentally regulated E2-E3 ubiquitylation modules, which couples remodelling of multi-protein E3 complexes with cognate E2 availability. This mechanism assures assembly of distinct erythroid maturation stage-dependent UBE2H-CTLH modules, required for the orderly progression of human erythropoiesis, thus establishing a paradigm for other E2-E3 modules involved in development processes.

## MATERIALS AND METHODS

### HUDEP2 and K562 cell culture and manipulation

HUDEP2 cells were cultured as described (Kurita *et al.*, 2013). Immature cells were expanded in StemSpan serum free medium (SFEM; Stem Cell Technologies) supplemented with 50 ng/ml human stem cell factor (hSCF) (R&D, ‘7466-SC-500), 3 IU/ml erythropoietin (EPO) (R&D, #287-TC-500), 1 μm dexamethasone (Sigma, #D4902), and 1μg/ml doxycycline (Sigma, #D3072). Cell densities were kept within 50×10^3^-0.8×10^6^ cells/ml and media replaced every other day. To induce erythroid maturation, HUDEP2 cells were cultured for three days (phase 1) in differentiation medium composed of IMDM base medium (GIBCO) supplemented with 2% (v/v) FCS, 3% (v/v) human serum AB-type, 3 IU/ml EPO, 10 μg/ml insulin, 3 U/ml heparin, 1 mg/ml holo-transferrin, 50 ng/ml hSCF, and 1 μg/ml doxycycline, followed by 9 days (phase 2) in differentiation medium without hSCF. Erythroid differentiation and maturation were monitored by flow cytometry (LSRFortessa, BD Biosciences), using PE-conjugated anti-CD235a/GYPA (BD Biosciences, clHIR2, #555570), FITC conjugated anti-CD49d (BD Biosciences, cl9F10, #304316), and APC-conjugated anti-Band3/SLC4A1 (gift from Xiulan An lab, New York Blood Centre) and analysed with FlowJo software.

The erythroleukemia cell lines K562 was obtained from ATCC (CCL-243^TM^) and cultured in IMDM (GIBCO) supplemented with 10% (v/v) FCS (GIBCO) and antibiotics (100 U/ml penicillin, 0.1 mg/ml streptomycin, GIBCO), and regularly checked for the absence of mycoplasma contamination. To induce erythroid-like differentiation, K562 cells were treated with 0.3 mM or 0.6 mM Na-butyrate (NaB, Millipore) for 24 hrs, and megakaryocyte-like differentiation was induced with 10 nM or 50 nM 12-O-Tetradodecanoyl-phorbol-13 acetate (TPA, Sigma) for 24 hrs. Where indicated, cells were treated with 10 μM proteasome inhibitor MG132 (Sigma) for different time phases.

### Plasmid preparation and mutagenesis

The cDNAs for MAEA and UBE2H, corresponding to the canonical UniProt sequences, were obtained from human cDNA library (Max-Planck-Institute of Biochemistry). 3xFlag- and 6xMyc-tagged constructs, using pcDNA5-FRT/TO as parental vector, were generated by classic recombinant cloning methods. Mutant versions of MAEA and UBE2H were prepared by the QuickChange protocol (Stratagene). All coding sequences were verified by DNA sequencing.

### K562 cell transfections, generation of CRISPR-Cas9 knock out cell lines

K562 cells were transformed by electroporation with Nucleofector Kit V (Bioscience, Lonza) according to the manufacturer’s protocol. Briefly, 1 × 10^6^ cell were harvested, washed once with 1xPBS (at room temperature), resuspended in 100μl Nucleofector solution, and mixed with 5 μg plasmid DNA. After electroporation, cells were recovered in 3 ml medium and cultured for 48 hrs. For immune precipitation (IP) experiments, three electroporation reactions with 1 × 10^6^ cells were done in parallel, transformed cells pooled and cultured for 48hrs.

*MAEA^-/-^, MKLN1^-/-^*, and *WDR26^-/-^* knock out cell lines were described previously (Sherpa *et al.*, 2021). To generate CRISPR-Cas9-(D10A) nickase-mediated functional knock outs of UBE2H, paired sense and anti-sense guide RNAs (gRNA) were designed to target exon 2 of UBE2H (Figure 4 **Supplemental figure 1B**). Sense and antisense gRNAs were cloned into pBABED-U6-Puromycine plasmid (gift from Thomas Macartney, University of Dundee, UK) and pX335-Cas9(D10a) (Addgene) (Cong *et al*, 2013), respectively. The plasmid pair was co-transfected into K562 cells using Lipofectamine LTX reagent (Invitrogene) according to the manufacturer’s protocol. Twenty-four hours after transfection, cells were selected in 2 μg/ml puromycin for two days, followed by expansion and single cell dilution to obtain cell clones. Successful knock out of UBE2H was validated by immunoblot analysis and genomic sequencing of the targeted locus (Figure 4A and figure 4 **Supplemental figure 1B**).

### Generation of CRISPR-Cas9-edited HUDEP2 knock out cell lines

To generate CRISPR-Cas9-(D10A) nickase-mediated functional knock outs of UBE2H in HUDEP2 cells, the same gRNA pair as described for the K562 knock out cell line has been used. For the functional knock outs of MAEA, RANBP9, and RANBP10 paired sense and anti-sense guide RNAs (gRNA) were designed to target exon 2 (MAEA), exon1 (RANBP9), and exon 7 (RANBP10) (Figure 2 **Supplemental figure 1 and** Figure 4 **Supplemental figure 1A**). The plasmid pairs were co-electroporated into HUDEP2 cells using Nucleofector Kit CD34^+^ (Bioscience, Lonza) according to the manufacturer’s protocol. Twenty-four hours after transfection, cells were selected in 2 μg/ml puromycin for two days, followed by expansion and single cell dilution to obtain cell clones. Cell densities were kept below 0.6 × 10^6^ cells/ml throughout the process. Successful knock outs were validated by immunoblot analysis and genomic sequencing of targeted loci (Figure 2C and 4D, Figure 2 **Supplemental figure 1 and** Figure 4 **Supplemental figure 1A and 1B**).

### Cell lysate preparation, immunoblot analysis, fractionation by sucrose density gradient, immunoprecipitation

To generate K562 and HUDEP2 cell lysates, cells were harvested by centrifugation at 360 × g, washed once with ice-cold 1xPBS, and resuspended in lysis buffer (40 mM HEPES pH7.5, 120 mM NaCl, 1mM EGTA, 0.5% NP40, 1mM DTT, and Complete protease inhibitor mix (Roche), and incubated on ice for 10 min. Cells were homogenized by pushing them ten times through a 23G syringe. The obtained lysates were cleared by centrifugation at 23,000 × g for 30 min at 4°C, and protein concentration was determined by Micro BCA-Protein Assay (Thermo Scientific, # 23235).

For immunoblot analysis lysates were denatured with SDS sample buffer, boiled at 95°C for 5 min, separated on SDS PAGE, and proteins were visualized by immunoblotting using indicated primary antibodies: RMND5a (Santa Cruz), MAEA (R&D systems), RANBP9 (Novus Biologicals), RANBP10 (Invitrogen, #PA5-110267), TWA1 (Thermo Fisher), ARMC8 (Santa Cruz), WDR26 (Bethyl Laboratories), MKLN1 (Santa Cruz), YPEL5 (Thermo Fisher), GID4 (described in (Sherpa *et al.*, 2021)), CD235a/GYPA (Abcam), HBD (Cell-Signaling), HBG1/2 (Cell-Signaling), and Flag (Sigma). Antibodies that recognize UBE2H were generated by immunizing sheep with GST-UBE2H (full length). Blots were developed using Clarity Western ECL Substrate (BioRad) and imaged using Amersham Imager 600 (GE Lifesciences). For quantitation described in Figure 6H, immunoblots from at least three biological repetitions were scanned with an Amersham Biosciences Imager 600 (GE Healthcare) and analyzed using ImageJ software.

For the sucrose gradient fractionation, 3 mg of total protein were loaded on to of a continues 5%-40% sucrose gradient (weight/volume in 40 mM HEPES pH7.5, 120 mM NaCl, 1mM EGTA, 0.5% NP40, 1mM DTT, and Complete protease inhibitor mix (Roche)) and centrifuged in a SW60 rotor at 34,300 rpm for 16 hours at 4°C. Thirteen 300 μl fractions were collected from top of the gradient, separated by SDS-PAGE, followed by immunoblotting using indicated antibodies.

Flag-tagged proteins were captured from 1 mg total cell lysate using anti-Flag affinity matrix (Sigma) for 1 hour at 4°C. For immunoprecipitation of endogenous proteins, 50 μg of antibody (ARMC8 or UBE2H) was incubated overnight with 4 mg of cell lysate at 4°C. In all, 30 μl of Protein-G agarose (Sigma) was added and the reaction was incubated for a further 2 hours. All immunoprecipitation reactions were washed in lysis buffer to remove non-specific binding, immunoadsorbed proteins eluted by boiling in reducing SDS sample buffer, separated by SDS-PAGE followed by immunoblotting using indicated antibodies.

### Nanobody production

Phage display selections: Purified human RANBP9-CTLH complex was coated on 96-well MaxiSorp plates by adding 100 μL of 1 μM proteins and incubating overnight at 4°C. Five rounds of phage display selections were then performed following standard protocols (Tonikian *et al*, 2007). The phage-displayed VH library used was reported before (Nilvebrant & Sidhu, 2018). Individual phage with improved binding properties obtained from round 4 and round 5 are identified by phage ELISA and subjected to DNA sequencing of the phagemids to obtain VH sequences. Phage ELISA with immobilized proteins were performed as described before (Zhang *et al*, 2016).

Cloning and protein purification: The nanobody cDNA was cloned into a vector containing either a C-terminal His tag (used for cryo-EM experiments) or an N-terminal GST tag (used for *in vivo* pulldown assays). The nanobody expression was carried out using BL21 pRIL cells and was purified from *E. coli* using either a Ni-NTA or a glutathione affinity chromatography, followed by size exclusion chromatography (SEC) in the final buffer containing 25 mM HEPES pH 7.5, 200 mM NaCl and 1 mM DTT.

### Determination of heme in cell lysates

HUDEP2 cell lines in proliferation media were allowed to grow to a maximum cell density of 0.8×10^6^ / ml in 12-Well plates. Cells were counted by an automated cell counter (Luna II, Logos Biosystems) before they were collected by centrifugation for 3 min, 300 × g. Cell pellets of 1.5×10^6^ cells were flash frozen in liquid nitrogen and stored at −80°C until further use. Cells were lysed in ice-cold HEPES-Triton-Lysis buffer (40 mM HEPES pH 7.5, 120 mM NaCl, 1 mM EDTA, 1 % Triton X-100, and cOmplete protease inhibitor (Roche)) with 1 μl of Lysis buffer per 10^4^ cells and kept on ice for 5 min. Cell lysates were cleared by centrifugation for 20 min, at 21000 × g at 4 °C and carefully transferred into new tubes. A second centrifugation step and transfer was performed to remove lipid residues. Protein concentration was determined by Micro BCA-Protein Assay (Thermo Scientific, # 23235). Lysate samples were diluted to 1 μg/μl with DPBS (GIBCO) and spectra analyses were done with 1.5 μl on a Nanodrop Spectrophotometer (Implen, NP80). The wavelength scans with peak maxima around 414 nm correspond to globin bound heme in differentiated HUDEP2 cell supernatants (Ghosh *et al*, 2018).

### RNA isolation and RT-qPCR

Cells were treated with indicated concentrations of Sodium butyrate (NaB) and TPA, respectively. 10×10^6^ cells were lysed in 1 ml Trizol (ThermoScientific, #15596018). 200 μL chloroform (FisherChemical, C496017) was added and samples were vigorously mixed. For phase separation, samples were then centrifuged at 10000x g for 10 min at 4°C. Subsequently, 400 μl of the upper clear phase were transferred into a new tube containing 500 μL isopropanol, mixed and incubated for 30min on ice, followed by centrifugation at 10000x g at 4°C for 10 mins. Pellet was washed once with 500 μl 70% ethanol and resuspended in RNase free water (Invitrogen, 10977-035). Samples were stored at −80°C until analysis. cDNA was generated using SuperScript IV First Strand Synthesis System (Invitrogen, 18091050) according to the manufacturer’s protocol. For qRT-PCR, cDNA, primers and SsoAdvanced Universal SYBR Green Supermix (BioRad, 1725274) were mixed and run on a CFX96 Touch Deep Well Real Time PCR System (Biorad) as per manufacturer’s protocol. Following forward/reverse primer pairs were used: for GAPDH 5’ GTTCGACAGTCAGCCGCATC / 5’ GGAATTTGCCATGGGTGGA; UBE2H 5’ CCTTCCTGCCTCAGTTATTGGC / 5’ CCGTGGCGTATTTCTGGATGTAC; GYPA 5’ ATATGCAGCCACTCCTAGAGCTC / 5’ CTGGTTCAGAGAAATGATGGGCA. Data was analyzed with BioRad CFX Manager using GAPDH for normalization.

### Protein expression and purification

The human RANBP10-and RANBP9-CTLH (TWA1-ARMC8-MAEA-RMND5A with either RANBP10 or RANBP9) complexes were purified from insect cell lysates using StrepTactin affinity chromatography, followed by anion exchange chromatography and size exclusion chromatography (SEC) in the final buffer containing 25 mM HEPES pH 7.5, 200 mM NaCl and 5 mM (for Cryo EM) or 1 mM DTT (for biochemical assays). N terminal GST-tagged version of hGid4(Δ1-99) and UBE2H were expressed in bacteria and purified by glutathione affinity chromatography followed by overnight cleavage using tobacco etch virus (TEV) protease. Further purification was carried out by anion exchange chromatography followed by size exclusion chromatography (SEC) in the final buffer containing 25 mM HEPES pH 7.5, 200 mM NaCl and 1 mM DTT. To obtain saturated RANBP10-CTLH complex with hGid4 for cryo EM analysis, it was added in 2-fold excess to TWA1-ARMC8-MAEA-RMND5A-RANBP10 before final SEC.

Untagged WT ubiquitin used for *in vitro* assays was purified via glacial acetic acid method (Kaiser *et al*, 2011), followed by gravity S column ion exchange chromatography and SEC.

### Ubiquitylation assay

The *in vitro* multi-turnover ubiquitylation assays with RANPB9-CTLH or RANBP10-CTLH complexes were performed using a C-terminally fluorescent-tagged model peptide with an N-terminal hGID4-interacting sequence PGLW and a single lysine placed at the 23^rd^ or 27^th^ position from the N terminus. The reaction was started by mixing 0.2 μM Uba1, 1 μM Ube2H, 0.5 μM RANBP9-TWA1-ARMC8-RMND5A-MAEA or RANBP10-TWA1-ARMC8-RMND5A-MAEA complex,1 μM hGid4, 1 μM fluorescent model peptide substrate and 20 μM Ub. The reaction was quenched in sample loading buffer at different timepoints and visualized by scanning the SDS-PAGE in the Typhoon Imager (GE Healthcare).

### Analytical SEC for RANBP10- and RANBP9-CTLH complexes

To see if the RANBP10 and RANBP9 complexes assemble and migrate at similar molecular weight range, analytical size exclusion chromatography was performed in a Superose 6 column (GE Healthcare) which was fitted to the Thermo Scientific Vanquish HPLC system. The column was equilibrated with 25 mM Hepes 7.5, 150 mM NaCl and 5 mM DTT and a 60 µl each of 10 µM purified RANBP10/TWA1/ARMC8/hGid4/RMND5A/MAEA and RANBP9/TWA1/ARMC8/hGid4/RMND5A/MAEA complexes were run through the HPLC system consecutively. The SEC fractions obtained were analyzed with Coomassie-stained SDS-PAGE.

### Cryo EM sample preparation and processing

Cryo EM grids were prepared using Vitrobot Mark IV (Thermo Fisher Scientific) maintained at 4°C and 100% humidity. 3.5 μl of the purified RANBP10-CTLH (RANBP9-TWA1-ARMC8-RMND5A-MAEA-hGid4) complex at 0.6 mg/ml was applied to Quantifoil holey carbon grids (R1.2/1.3 300 mesh) that were glow-discharged separately in Plasma Cleaner. After sample application grids were blotted with Whatman no. 1 filter paper (blot time: 3 s, blot force: 3) and vitrified by plunging into liquid ethane.

For the nanobody bound RANBP9-CTLH complex, the purified nanobody was first mixed to the RANBP9-CTLH complex and ran on size exclusion chromatography. The peak fraction was then concentrated and prepared for Cryo-EM using the same approach as mentioned above.

Both the Cryo EM data were collected on a Talos Arctica transmission electron microscope (Thermo Fisher Scientific) operated at 200 kV, equipped with a Falcon III (Thermo Fisher Scientific) direct electron detector, respectively. The data collection was carried out using EPU software (Thermo Fisher Scientific).

The Cryo EM data processing was carried out with Relion (Fernandez-Leiro & Scheres, 2017; Scheres, 2012; Zivanov *et al*, 2018). For processing the micrographs, frames were first motion-corrected using Relion’s own implementation of MotionCor like algorithm by Takanori Nakane followed by contrast transfer function estimation using CTFFind 4.1. Particles were auto-picked using Gautomatch (http://www.mrc-lmb.cam.ac.uk/kzhang/) using a template of RANBP9-CTLH^SR4^ (EMDB: EMD-12537). The extracted particles were subjected to several rounds of 2D classification and 3D classification followed by auto-refinement without and with a mask. To improve the quality of maps obtained for RANBP10-CTLH complex, a focused 3D classification without particle alignment was performed with a mask over CTLH^SRS^. The best class with most features were chosen and auto-refinement with mask over the CTLH^SRS^ was performed followed by post-processing.

### MS-based proteomics analysis of HUDEP2 samples

Cell pellets were lysed in SDC buffer (1% sodium deoxycholate in100 mM Tris pH 8.5) and then heated for 5 min at 95°C. Lysates were cooled on ice and sonicated for 15 min at 4°. Protein concentration was determined by Tryptophan assay as described in (Kulak *et al*, 2014) and equal amount of proteins were reduced and alkylated by 10 mM TCEP and 40 mM 2-Chloroacetamide, respectively, for 5 min at 45°C. Proteins were subsequently digested by the addition of 1:100 LysC and Trypsin overnight at 37°C with agitation (1,500 rpm). Next day, around 10 μg of protein material was processed using an in-StageTip (iST) protocol (Kulak *et al.*, 2014). Briefly, samples were at least 4-fold diluted with 1% trifluoro-acetic acid (TFA) in isopropanol to a final volume of 200 μL and loaded onto SDB-RPS StageTips (Empore). Tips are then washed with 200μL of 1% TFA in isopropanol and 200μL 0.2% TFA/2% ACN (acetonitrile). Peptides were eluted with 80 μl of 1.25% Ammonium hydroxide (NH4OH)/80% ACN and dried using a SpeedVac centrifuge (Concentrator Plus; Eppendorf). MS loading buffer (0.2% TFA/2%ACN (v/v)) was added to the dried samples prior to LC-MS/MS analysis. Peptide concentrations were measured optically at 280 nm (Nanodrop 2000; Thermo Scientific) and subsequently equalized using MS loading buffer. Approximately, 300-500 ng peptide from each sample was analyzed using a 100 min gradient single shot DIA method.

### LC-MS/MS analysis and data processing

Nanoflow LC-MS/MS measurements were carried out on an EASY-nLC 1200 system (Thermo Fisher Scientific) coupled to the Orbitrap instrument, namely Q Exactive HF-X and a nano-electrospray ion source (Thermo Fisher Scientific). We used a 50 cm HPLC column (75 μm inner diameter, in-house packed into the tip with ReproSil-Pur C18-AQ1.9 μm resin (Dr. Maisch GmbH)). Column temperature was kept at 60°C with an in-house developed oven.

Peptides were loaded in buffer A (0.1 % formic acid (FA) (v/v)) and eluted with a linear 80 min gradient of 5–30 % of buffer B (80 % acetonitrile (ACN) and 0.1% FA (v/v)), followed by a 4 min increase to 60 % of buffer B and a 4 min increase to 95 % of buffer B, and a 4 min wash of 95% buffer B at a flow rate of 300 nl/min. Buffer B concentration was decreased to 4% in 4 min and stayed at 4% for 4 min. MS data were acquired using the MaxQuant Live software and a data-independent acquisition (DIA) mode (Wichmann *et al*, 2019). Full MS scans were acquired in the range of m/z 300-1,650 at a resolution of 60,000 at m/z 200 and the automatic gain control (AGC) set to 3e6. Full MS events were followed by 33 MS/MS windows per cycle at a resolution of 30,000 at m/z 200 and ions were accumulated to reach an AGC target value of 3e6 or an Xcalibur-automated maximum injection time. The spectra were recorded in profile mode.

The single shot DIA runs of HUDEP2 samples were searched with direct DIA (dDIA) mode in Spectronaut version 14 (Biognosys AG) for final protein identification and quantification. All searches were performed against the human SwissProt reference proteome of canonical and isoform sequences with 42,431 entries downloaded in July 2019. Carbamidomethylation was set as fixed modification and acetylation of the protein N-terminus and oxidation of methionine as variable modifications. Trypsin/P proteolytic cleavage rule was used with a maximum of two miscleavages permitted and a peptide length of 7–52 amino acids. A protein and precursor FDR of 1% were used for filtering and subsequent reporting in samples (q-value mode).

### Bioinformatics data analysis

We mainly performed data analysis in the Perseus (version 1.6.0.9 and 1.6.1.3) (Tyanova *et al*, 2016). Protein intensities were log2-transformed for further analysis. Data sets were filtered to make sure that identified proteins showed expression in all biological triplicates of at least one experimental group and the missing values were subsequently replaced by random numbers that were drawn from a normal distribution (width = 0.3 and down shift = 1.8). PCA of experimental groups and biological replicates was performed using Perseus. Multi-sample test (ANOVA) for determining if any of the means of experimental group was significantly different from each other was applied to protein data set. For truncation, we used permutation-based FDR which was set to 0.01 or 0.05 in conjunction with an S0-parameter of 0.1. For hierarchical clustering of significant proteins, median protein abundances of biological replicates were z-scored and clustered using Euclidean as a distance measure for row and/or column clustering. Gene Ontology (GO) enrichments in the clusters were calculated by Fisher’s exact test using Benjamini-Hochberg FDR for truncation. Mean log2 ratios of biological triplicates and the corresponding P-values were visualized with volcano plots and significance was based on a FDR < 0.05 or 0.01. Network representation of significantly regulated proteins was performed with the STRING app (1.5.1) in Cytoscape (3.7.2).

## ACKNOWLEDGEMENTS

This work was supported by the Max Planck Society for the Advancement of Science and by the Deutsche Forschungsgemeinschaft (DFG, German Research Foundation) – SCHU 3196/1-1. We thank all the members of the department of Molecular Machines and Signaling at Max Planck Institute of Biochemistry for their assistance and helpful discussions, especially J. Rajan Prabu for guidance in structural analysis, Susanne von Gronau for maintaining the insect cells and Josef Kellermann for maintaining the lab. We also thank Daniel Bollschweiler and Tillman Schäfer for maintaining the MPIB Cryo EM facility; Stephan Übel and Stefan Pettera in the MPIB biochemistry core facility for the peptide synthesis.

