## Supplemental Figures for "Differential UBE2H-CTLH E2-E3 ubiquitylation modules regulate erythroid maturation"

**SUPPLEMENTAL MATERIAL**

**Figure 1 Supplemental figure 1**
**Figure 2 Supplemental figure 1**
**Figure 2 Supplemental figure 2**
**Figure 3 Supplemental figure 1**
**Figure 3 Supplemental figure 2**
**Figure 4 Supplemental figure 1**
**Figure 4 Supplemental figure 2**
**Figure 5 Supplemental figure 1**
**Figure 6 Supplemental figure 1**

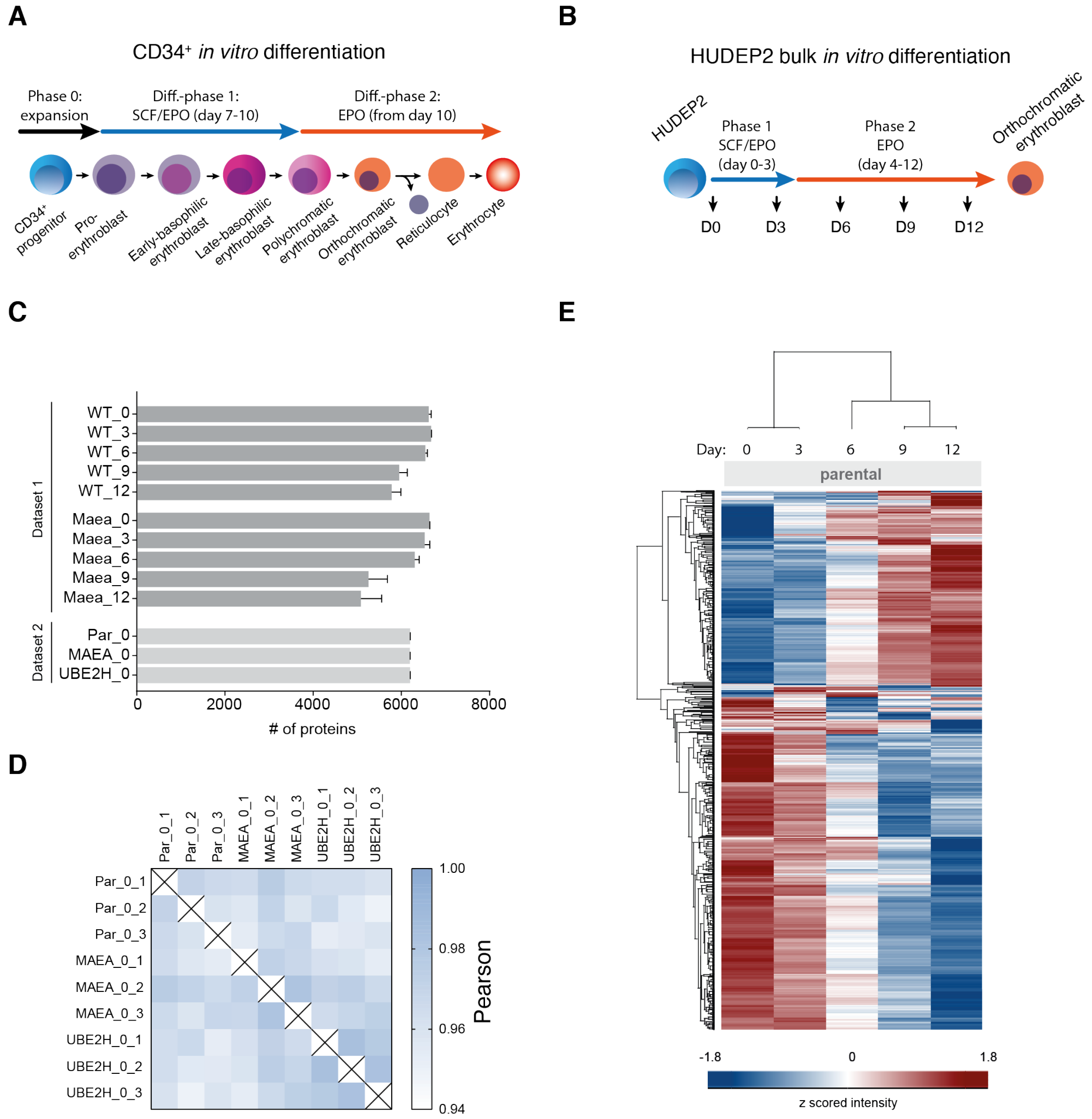


**Figure 1 Supplemental figure 1. A)** Schematic with 2-phase culture conditions of *in vitro* reconstituted erythropoiesis of CD34$^{+}$ cells. Stem cell factor, SCF; erythropoietin, EPR. **B)** Schematic with culture conditions of *in vitro* reconstituted erythropoiesis of HUDEP2 cells. **C)** Number of different proteins quantified in each differentiation stage (day 0, 3, 6, 9, 12) of HUDEP2 parental (WT) and MAEA$^{-/-}$cl3-1 cells (Dataset 1), and HUDEP2 parental, MAEA$^{-/-}$cl3-1, and UBE2H$^{-/-}$cl13 (Dataset 2). The mean value ± SD of three biological replicates are shown. **D)** Correlation based analysis illustrating reproducibility between biological replicates. High (1.0) and lower (0.94) Pearson correlation are indicated in blue and white, respectively. **E)** Heat map of z-scored protein abundance (log$_2$ DIA intensity) of 2771 differentially expressed proteins in differentiated HUDEP2 cells (ANOVA, FDR <0.01). Differentiation time stages correspond to schematic in B.

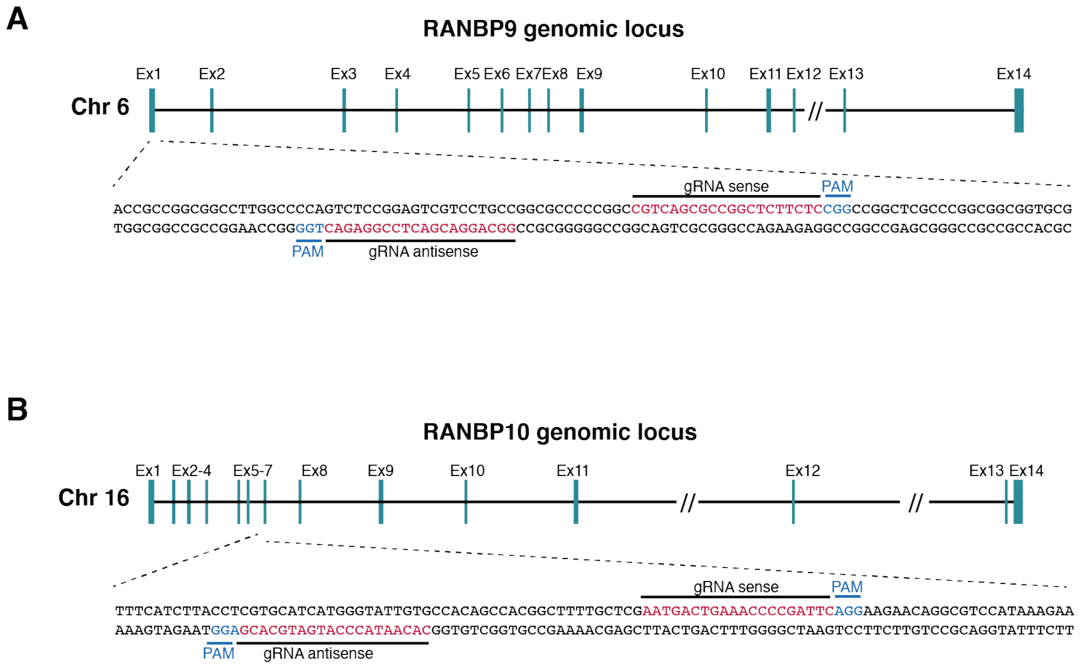


**Figure 2 Supplemental figure 1. A)** Schematics of CRISPR knockout of RANBP9, showing exon structure and indicated guide RNAs and PAM sequences. **B)** Schematics of CRISPR knockout of RANBP10, showing exon structure and indicated guide RNAs and PAM sequences.

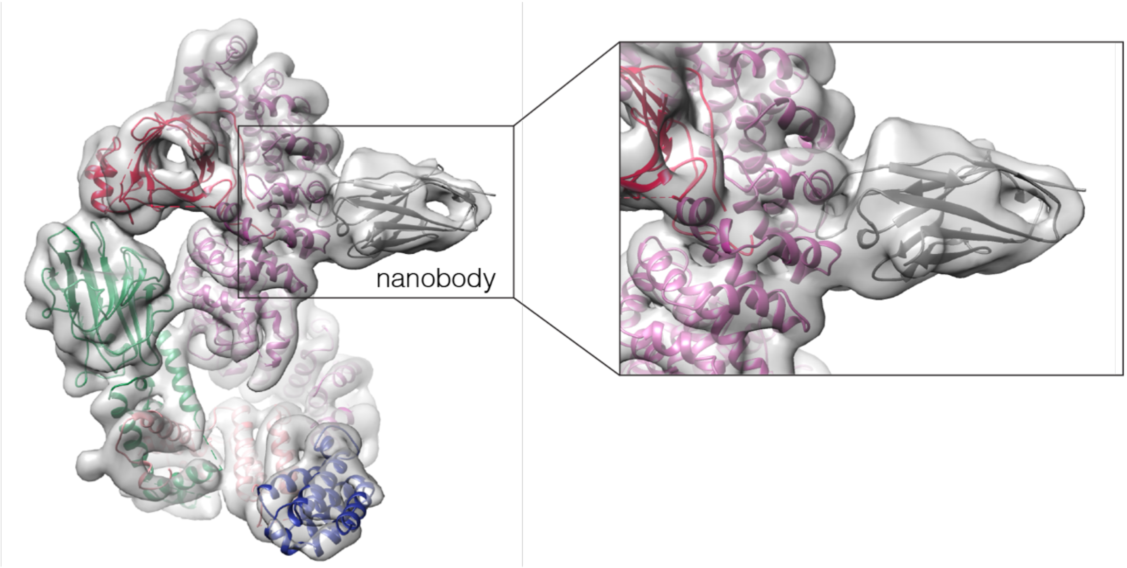


**Figure 2 Supplemental figure 2.** Cryo EM structure of the ARMC8-specific nanobody bound to CTLH core. The homology model of the nanobody (grey) and the structure of the CTLH core (color-coded by subunits as in Figure 3C, PDB: 7NSC) was fitted in the cryo EM density (transparent surface). The CDR loop of the nanobody contacts the C-terminal Armadillo repeats of ARMC8 (purple) ranging from aa 420 to 470.

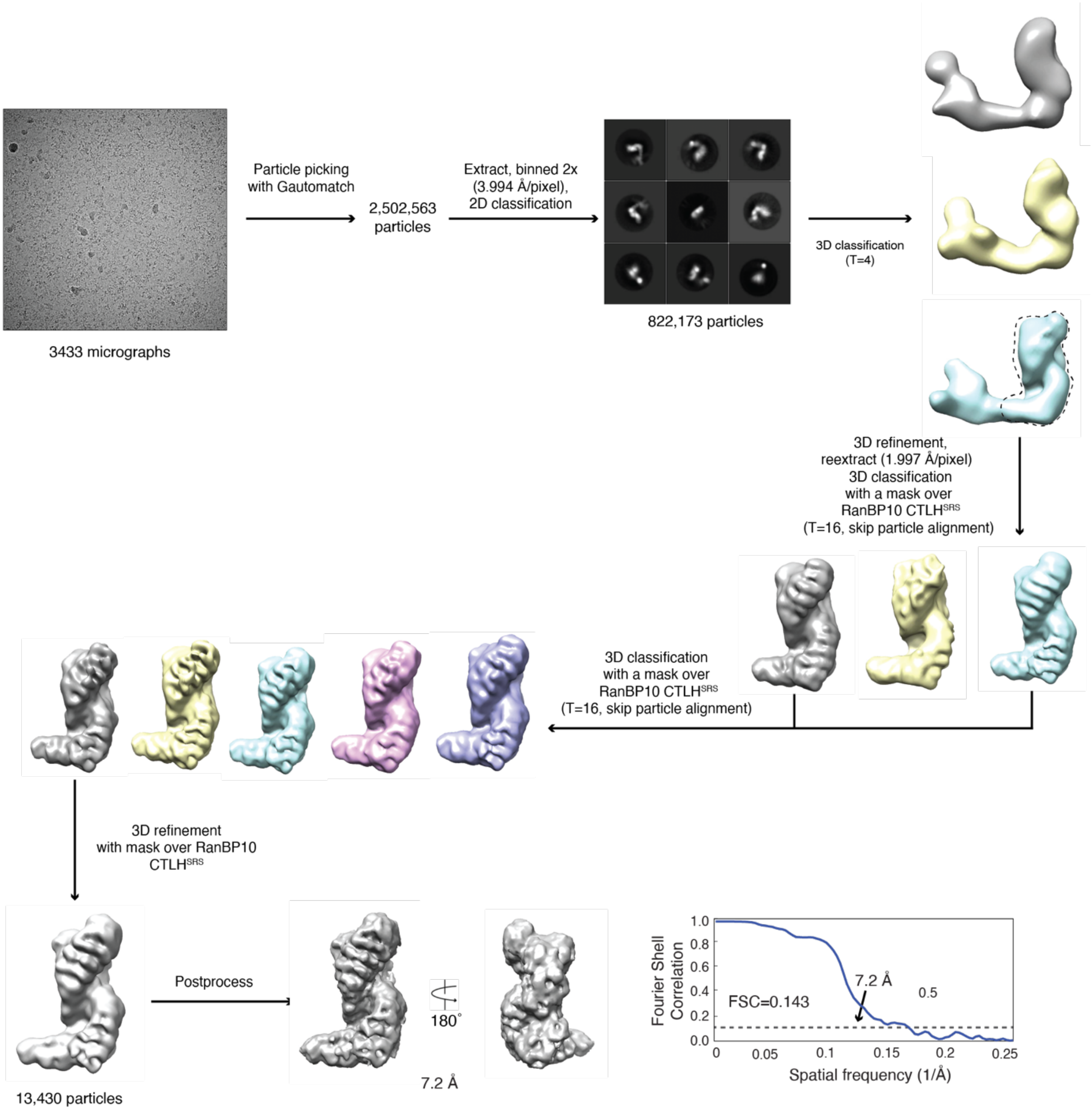


**Figure 3 Supplemental figure 1.** Flowchart of cryo EM processing for the RANBP10-CTLH complex dataset.

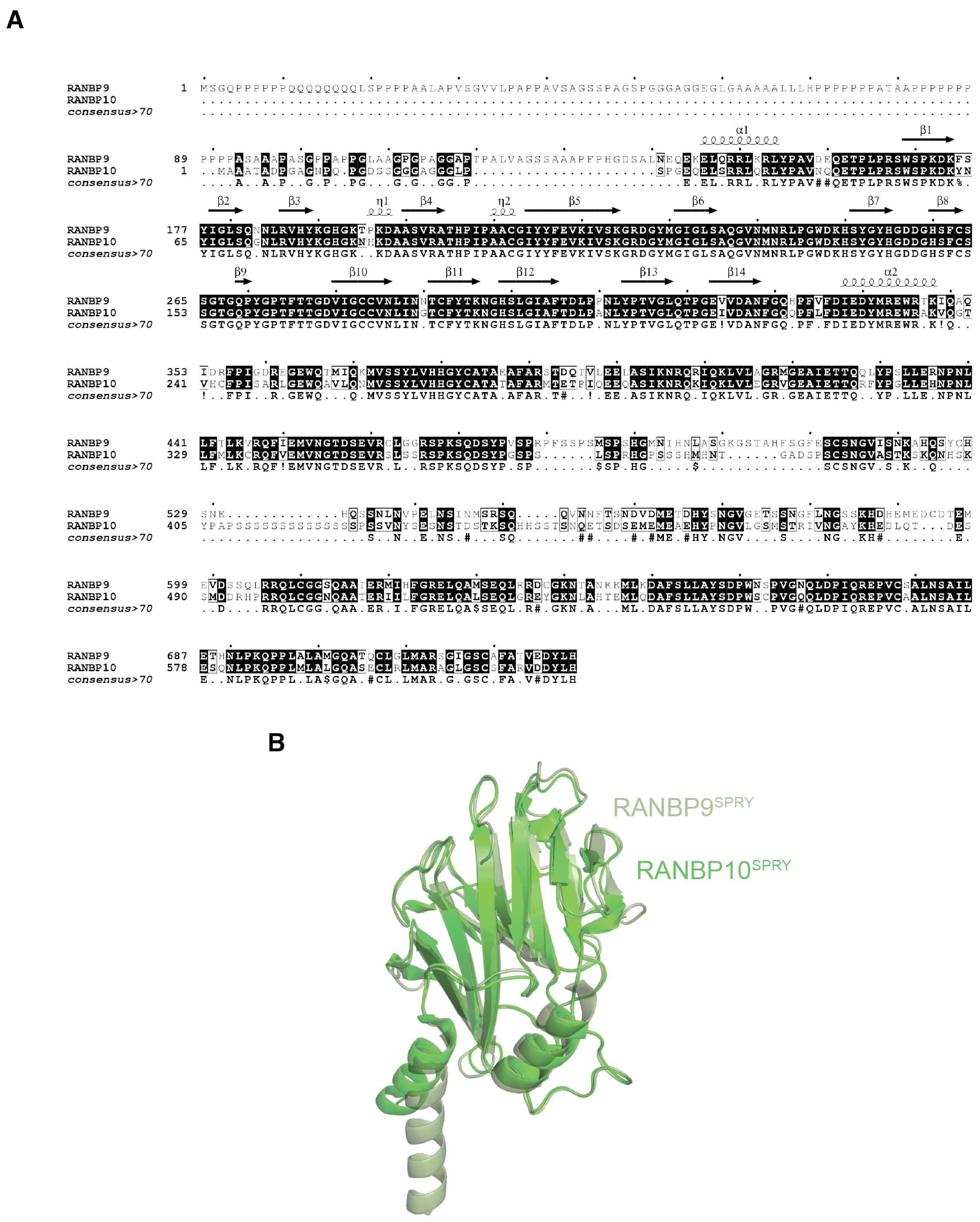


**Figure 3 Supplemental figure 2. A)** Alignment of human RANBP9 and RANBP10 sequences, with identical sequences highlighted in black. **B)** Structural alignments of SPRY domains of RANBP9 (light green) and RANBP10 (dark green).

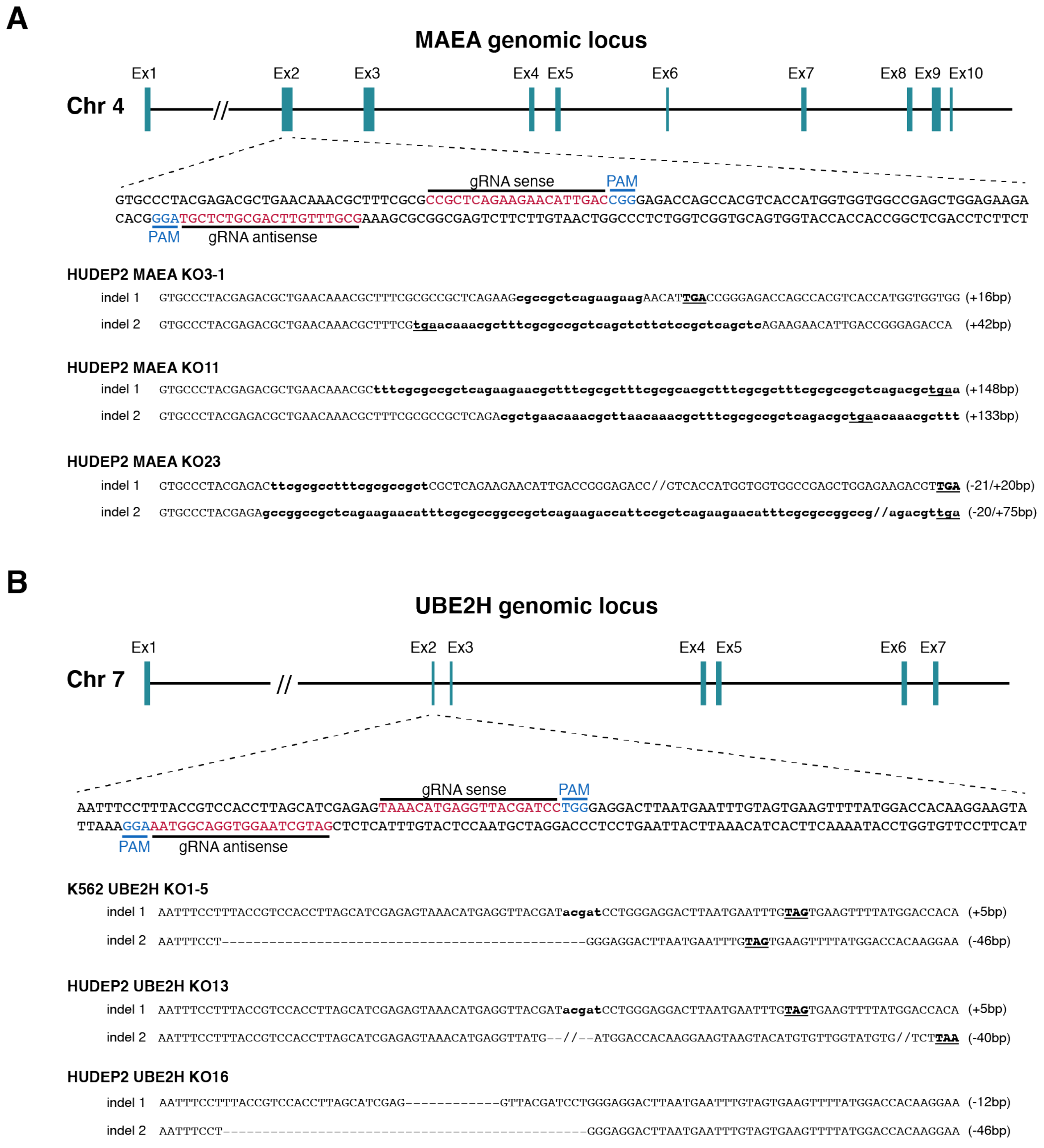


**Figure 4 Supplemental figure 1. A)** Schematics of CRISPR knockouts of MAEA, showing exon structure and indicated guide RNAs and PAM sequences. Insertions/deletions (indels) of each HUDEP2 MAEA knockout clones are presented. **B)** Schematics of CRISPR knockouts of UBE2H, showing exon structure and indicated guide RNAs and PAM sequences. Insertions/deletions (indels) of K562 and each HUDEP2 UBE2H knockout clones are presented.

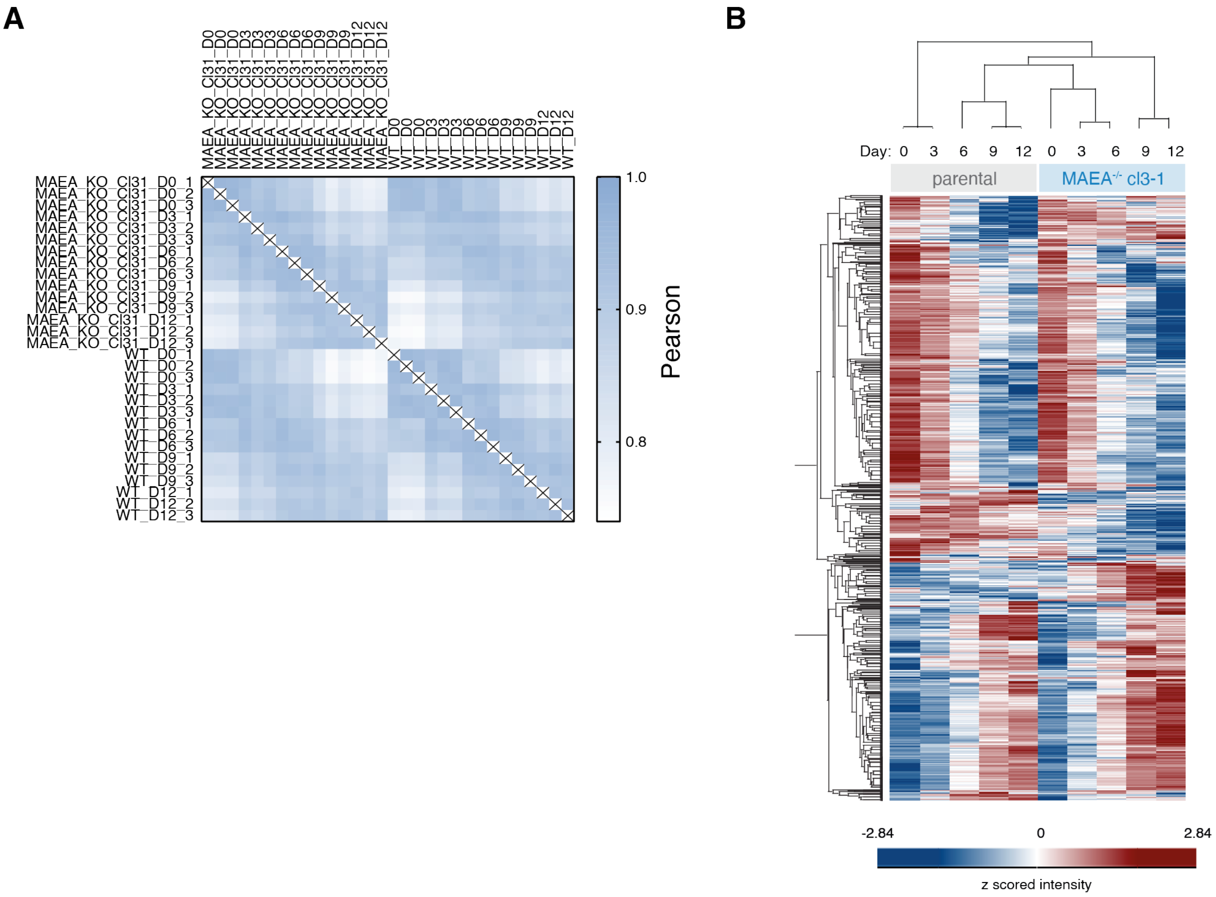


**Figure 4 Supplemental figure 2. A)** Correlation based analysis illustrating reproducibility between biological replicates. High (1.0) and lower (0.75) Pearson correlation are indicated in blue and white, respectively. **B)** Heat map of z-scored protein abundance ($log_2$ DIA intensity) of 3193 differentially expressed proteins in differentiated HUDEP2 parental and MAEA$^{-/-}$cl3-1 cells (ANOVA, FDR <0.01).

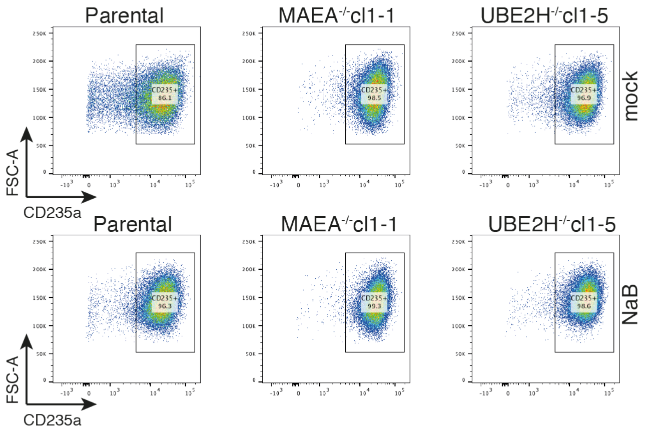


**Figure 5 Supplemental figure 1.** Flow cytometry blots of mock (top) or NaB-treated (bottom) K562 parental, MAEA$^{-/-}$cl1-1 and UBE2H$^{-/-}$cl1-5 cell lines showing CD235a expression.

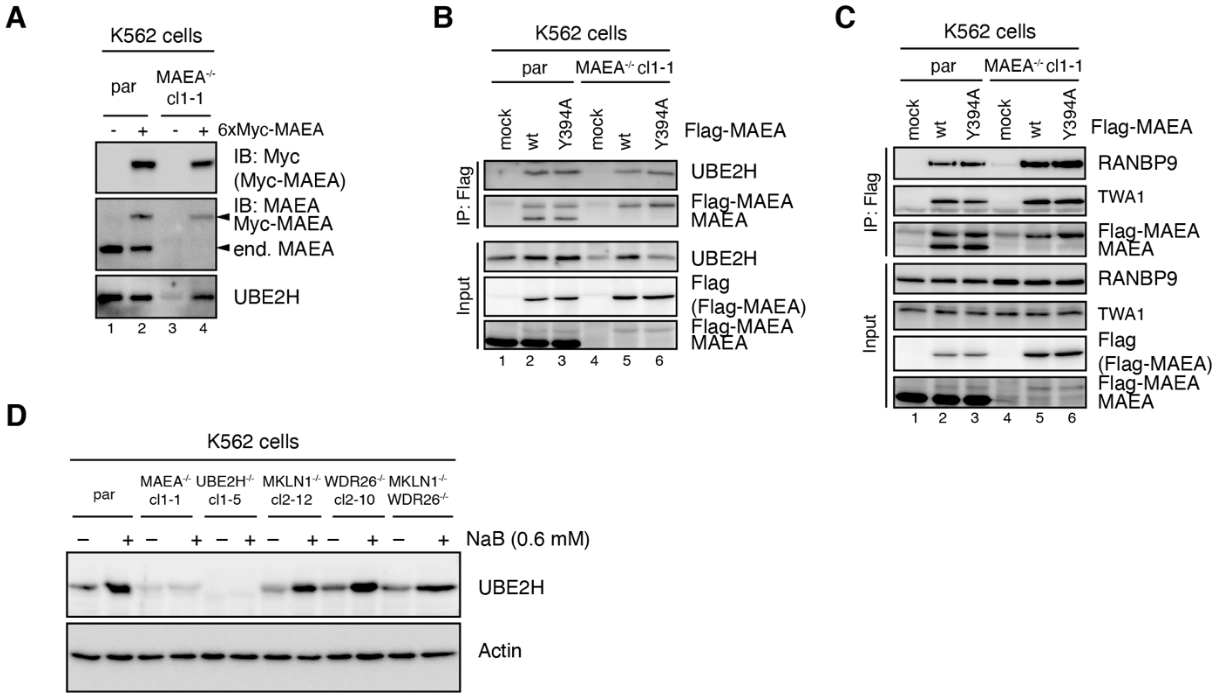


**Figure 6 Supplemental figure 1. A)** K562 parental and MAEA$^{-/-}$cl1-1 cells were mock (-) or with 6xMyc-tagged MAEA (+) transfected and cell lysates analysed for UBE2H protein levels by immunoblot analysis. **B)** and **C)** K562 parental and MAEA$^{-/-}$cl1-1 cells were mock, Flag-MAEA wildtype (WT), or mutated Flag-MAEA-Y394A (Y394A) transfected and Flag-IPs from cell lysates were analysed by immunoblotting using indicated antibodies. **D)** K562 parental and knock out cell lines were treated with 0.6 mM NaB for 24 hours and analysed for UBE2H protein levels by immunoblotting.
